# Neurodevelopmentally rooted epicenters in schizophrenia: sensorimotor-association spatial axis of cortical thickness alterations

**DOI:** 10.1101/2024.03.13.584752

**Authors:** Yun-Shuang Fan, Yong Xu, Meike Dorothee Hettwer, Pengfei Yang, Wei Sheng, Chong Wang, Mi Yang, Matthias Kirschner, Sofie Louise Valk, Huafu Chen

**Affiliations:** The Clinical Hospital of Chengdu Brain Science Institute, School of Life Science and Technology, University of Electronic Science and Technology of China, Chengdu, China; Otto Hahn Group Cognitive Neurogenetics, Max Planck Institute for Human Cognitive and Brain Sciences, Leipzig, Germany; Department of Psychiatry, First Hospital/First Clinical Medical College of Shanxi Medical University, Taiyuan, China; MOE Key Lab for Neuroinformation, High-Field Magnetic Resonance Brain Imaging Key Laboratory of Sichuan Province, University of Electronic Science and Technology of China, Chengdu, China; Institute of Neuroscience and Medicine (INM-7: Brain and Behavior), Research Centre Jülich, Jülich, Germany; Max Planck School of Cognition, Leipzig, Germany; Institute of Systems Neuroscience, Medical Faculty, Heinrich Heine University Düsseldorf, Düsseldorf, Germany; Division of Adult Psychiatry, Department of Psychiatry, University Hospitals of Geneva, Geneva, Switzerland

**Keywords:** connectome, cortical thickness, early-onset schizophrenia, neurodevelopment, transcriptomics

## Abstract

Pathologic perturbations in schizophrenia have been suggested to propagate via the functional and structural connectome across the lifespan. Yet how the connectome guides early cortical reorganization of developing schizophrenia remains unknown. Here, we used early-onset schizophrenia (EOS) as a neurodevelopmental disease model to investigate putative early pathologic origins that propagate through the functional and structural connectome. We compared 95 patients with antipsychotic-naïve first-episode EOS and 99 typically developing controls (7–17 years of age, 120 females). Whereas patients showed widespread cortical thickness reductions, thickness increases were observed in primary cortical areas. Using normative connectomics models, we found that epicenters of thickness reductions were situated in association regions linked to language, affective, and cognitive functions, while epicenters of increased thickness in EOS were located in sensorimotor regions subserving visual, somatosensory, and motor functions. Using post-mortem transcriptomic data of six donors, we observed that the epicenter map differentiated oligodendrocyte-related transcriptional changes at its sensory apex and the association end was related to expression of excitatory/inhibitory neurons. More generally, we observed that the epicenter map was associated with neurodevelopmental disease gene dysregulation and human accelerated region genes, suggesting potential shared genetic determinants across various neurodevelopmental disorders. Taken together, our results underscore the developmentally rooted pathologic origins of schizophrenia and their transcriptomic overlap with other neurodevelopmental diseases.

## Introduction

Schizophrenia is increasingly conceptualized as a neurodevelopmental disorder with a polygenic architecture (1, 2), in which pathologic processes originate early in brain development (3). However, why, when, and where these alterations occur in the brain is incompletely understood. During development, the human brain shows systematic patterns of maturation along anatomically and functionally connected regions (4), called the connectome. Despite the many biological and functional benefits for resource sharing through a refined connectome, pathologic perturbations have also been found to propagate via connections among regions in schizophrenia (5). Transmodal connectome has been reported to shape distributed deformation patterns in chronic schizophrenia (6), while unique architecture constrains early patterns in first-episode schizophrenia (7, 8). Given the developmental factor, we hypothesize that pathologic processes in early-onset schizophrenia (EOS) is shaped by the developing intrinsic brain organization.

Numerous neuroimaging studies have reported that schizophrenia is associated with pronounced brain structural alterations, typically with widespread cortical thinning. For example, schizophrenia patients have disease-specific and progressive cortical thinning in the frontal and temporal regions compared to healthy controls (9, 10). During adolescent maturation, patients with early-onset schizophrenia (EOS) have been shown to exhibit progressive reorganization of the cortex, which is dominated by the insula and occipital cortex (11). Specifically, EOS patients have increased cortical thickness thinning in the pre- and post-central, frontal, and temporal regions, and reduced cortical thickness thinning in the occipital cortex during development (12, 13). Recent evidence suggests that schizophrenia-related brain alteration topography is not randomly distributed but follows the intrinsic network organization of the human connectome (14–16). Indeed, although these structurally altered regions are distant from one other, the regions are strongly interconnected (15). These convergent findings of schizophrenia point towards pathologic origins that propagate though the brain connectome, herein referred to as disease epicenters.

Based on the assumption that the degree to which regions are similarly affected by pathology is associated with their connections, disease epicenters can be identified as those regions with connectivity profiles that closely resemble disease-related brain alteration patterns (17). This novel epicenter model has determined disease-specific epicenters of multiple neurodegenerative diseases, revealing functional and structural network architecture underlying their brain abnormalities (18–20). Moreover, transdiagnostic epicenters across psychiatric disorders indicated common prefrontal and temporal network anchors spreading psychopathologic effects (21), regardless of whether they are caused by illness or medication (8). Schizophrenia-related tissue volume alteration patterns have been reported to be circumscribed by the ventral attention network, with a disease epicenter in the anterior cingulate cortex (6). In agreement with this finding, transmodal epicenters emerged as shared epicenters across disease stages, while occipital and parietal epicenters were additionally found in early courses of adult schizophrenia (7). However, the neurodevelopmental roots of disease epicenters have not been established. For example, how brain functional and structural connectomes guide early cortical reorganization in EOS throughout childhood and adolescence is unknown.

From childhood to adolescence, cortical maturation has been reported to occur in a systematic manner that progresses from primary sensorimotor cortices to transmodal association cortices subserving executive, socioemotional, and mentalizing functions (22). The hierarchical unfolding of cortical development is supported by genetic processes (23). The neurodevelopmental hypothesis of schizophrenia suggests that during this critical period, genetic and environmental risk factors jointly disturb brain maturation (24). Furthermore, schizophrenia and other neurodevelopmental disorders, such as autism spectrum disorder, have recently been recognized as part of a common neurodevelopmental continuum (25). Specifically, schizophrenia and other neurodevelopmental disorders have a shared molecular etiology and considerable genetic overlap (26). Recently, a promising evolution hypothesis was proposed to explain the neurodevelopmental continuum. This hypothesis posits that these mental illnesses emerge as costly by-products of human evolution (27). For example, the human accelerate region (HAR) genes, i.e., the human-specific genes located in the accelerated diverging HARs between humans and chimpanzee ancestors (28), may harbor common genetic determinants shared across different psychiatric disorders (29). Notably, the availability of whole-brain gene expression atlases from the Allen Human Brain Atlas ([AHBA]; http://human.brain-map.org) microarray dataset (30) offers an unprecedented chance to bridge the brain connectome and microscale gene transcriptomes (31). Research combining neuroimaging and gene transcripts have suggested that disease-specific brain alterations are underpinned by brain expression of disease-relevant genes (32). Therefore, these advances have enabled us to investigate the microscale neurobiological mechanism underlying schizophrenia brain phenotypes, which aids in further elucidating the disease pathogenesis from a neurodevelopmental continuum perspective.

In the current study, we used EOS patients, 7–17 years of age, served as a neurodevelopmental disease model to investigate the putative early pathologic origins that propagate through the brain connectome. We first identified early disease epicenters of EOS by assessing the influence of functional and structural connectivity profiles on the spatial distribution of cortical thickness alterations in patients, as in previous studies (7, 17, 33). We then contextualized these observations within a micro-level transcriptomic architecture by applying partial least squares (PLS) analysis to disease epicenters and AHBA gene expression maps (30). We further examined the relationship between epicenter-associated gene weights and differential gene expression of multiple major psychiatric disorders (34), and the association between epicenter-related genes and HAR genes (35). Overall, we found that early disease epicenters with thickness reductions were in sensorimotor cortices, whereas epicenters with increased thickness were in association cortices. Distinct microscale molecular processes were detected behind epicenters of cortical thinning and thickening in patients with EOS. Epicenter-related gene expression was associated with genetic dysregulation of schizophrenia, autism spectrum disorder, and bipolar disorder. Moreover, epicenter-related genes overlapped with HAR genes harboring common genetic determinants across these disorders.

## Materials and Methods

### Participants and imaging data preprocessing

A total of 199 pediatric participants, 7–17 years of age, were recruited from the First Hospital of Shanxi Medical University, China. They comprised 99 drug-naïve, first-episode EOS patients and 100 typically developing (TD) controls. Details of the imaging protocol have been published elsewhere (36), and here we repeat for clarity. In brief, multimodal imaging data were acquired on a 3 Tesla Siemens MAGNETOM Verio scanner at the First Hospital of Shanxi Medical University. From this original sample, four patients were excluded due to incomplete scanning data, and one patient was excluded due to poor quality cortical parcellation. A final sample including 95 EOS patients and 99 demographically matched TD controls were further analyzed; detailed demographic data are included in **Table S1**. All T1-weighted data were preprocessed with FreeSurfer package (v7.1.0, http://surfer.nmr.mgh.harvard.edu/), including cortical segmentation and surface reconstruction. rs-fMRI data were preprocessed with the CBIG pipeline (https://github.com/ThomasYeoLab/CBIG) based on FSL (v5.0.9) and FreeSurfer (v7.1.0), which included removal of the first four volumes, slice-timing, motion correction, boundary-based registration to structural images, covariates regression, and bandpass filtering (0.01–0.08 Hz). DTI data were preprocessed with FSL (FMRIB Software Library v5.0.9, http://www.fmrib.ox.ac.uk/fsl) and the diffusion toolkit, including eddy current correction, diffusion tensor model estimation and whole-brain fiber tracking. Additional details about the participants, imaging data acquisition, cortical thickness estimation, and normative pediatric connectivity matrix construction are included in Supplement.

### Disease epicenter mapping

We calculated disease epicenters in EOS following published ENIGMA pipelines (https://enigma-toolbox.readthedocs.io/en/latest/) (7, 37). Specifically, we correlated normative pediatric functional and structural connectomes spatially with the cortical thickness alteration map in patients with EOS. We additionally conducted a robustness check by calculating disease epicenters using a normative adult connectome from the Human Connectome Project data (38). This epicenter mapping analysis generated one correlation coefficient for each region (herein referred to as epicenter values), representing the association between the connectivity profile and disease-related abnormality map. Regions with high absolute correlation coefficient values were identified as disease epicenters, in which positive (negative) coefficients reflect positive (negative) epicenters. The statistical significance of spatial correlations was assessed using spin permutation tests that account for spatial autocorrelation (*p_spin_* < 0.05, 10,000 times) (39). Spin test details are shown in Supplement. Specifically, a region could potentially be an epicenter if the following criteria were met: (i) strongly connected to other high-thinned regions or weakly connected to other low-thinned regions (negative epicenters); and (ii) strongly connected to other high-thickened regions or weakly connected to other low-thickened regions (positive epicenters). Additionally, we delineated the relevance between disease epicenters and neurodevelopment, functional systems, and cognitive functions. Detailed steps about neurodevelopment, functional systems, and cognitive embedding are included in Supplement.

### Transcriptomic genetic decoding

To examine transcriptomic expression underlying disease epicenters, we used high-resolution microarray gene expression data from six post mortem brains provided by the AHBA (30). Gene expression data were processed and mapped onto 400 cortical parcels (40) through the abagen toolbox (https://abagen.readthedocs.io/) (41), yielding a 400 × 15,631 matrix (regions × genes) of transcriptional levels. Next, we used PLS analysis (42) to decompose associations between gene expression (X_400 × 15631_) and epicenter maps (Y_400 × 2_) into orthogonal sets of latent variables with maximum covariance. To identify involving cell types and biological pathways, we further performed cell type deconvolution using cell-specific aggregate gene sets, as proposed by previous studies (43), and enrichment analysis using Metascape (https://metascape.org/gp/index.html#/main/step1) (44). Detailed steps about gene expression data processing, PLS analysis, and identification of involving cell types and biological pathways are shown in Supplement.

### Correlation with major brain disorders and HAR genes

To evaluate the relationship between loadings of the gene sets identified by PLS analysis and disease-specific gene dysregulation, we used postmortem brain tissue measurements of mRNA (false discovery rate [FDR]; *p_FDR_*< 0.05) in six brain disorders, including schizophrenia, autism spectrum disorder, bipolar disorder, major depressive disorder, alcohol abuse disorder (alcoholism), and inflammatory bowel disease (34). Furthermore, we determined whether the transcriptomic architecture of genes located in HARs of the genome underpin disease epicenter map using 2143 HAR genes defined by a previous study (35). Detailed steps are shown in Supplement.

## Results

First, vertex-wise cortical thickness were down-sampled based on Freesurfer segmentation into a 400-parcel cortical Schaefer parcellation atlas (40) and the group differences were evaluated between EOS patients and TD controls using two-sample *t*-tests with covariates, including age and gender (*p_FDR_* < 0.05). In agreement with previous findings (13), patients with EOS had widespread cortical thickness reductions relative to TD controls (**Figure 1**), predominantly in the dorsal attention network, limbic network, and default mode network (DMN) (**Table S2**). In addition, patients had increased cortical thickness in the left primary visual regions. Group comparison results were similar after including age squared as covariates (**Figure S1**).

**Figure 1.**
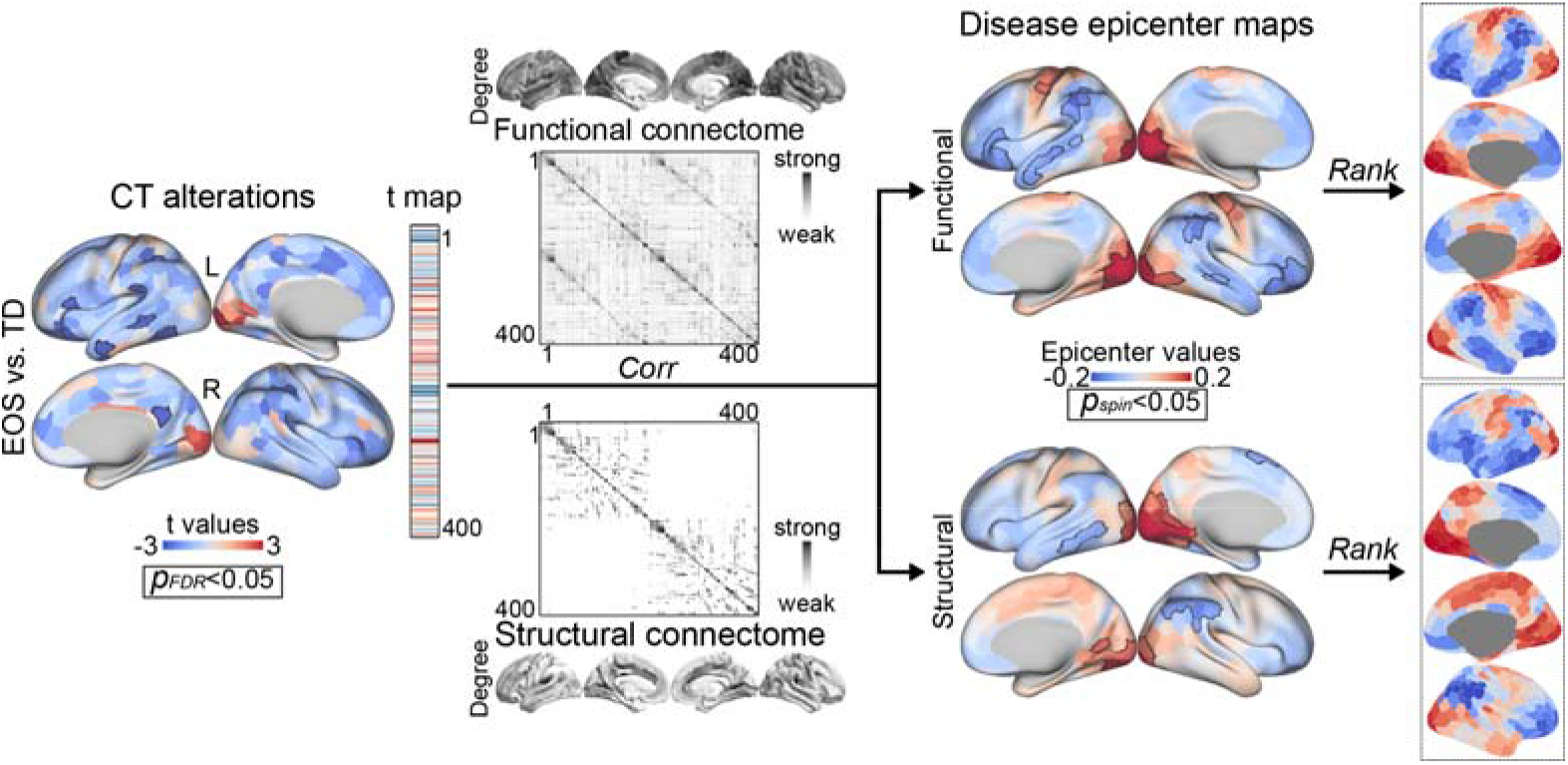
Disease epicenters for early-onset schizophrenia (EOS). Mapping disease epicenters by relating cortical thickness (CT) alterations and normative pediatric brain connectome. *Left:* cortical parcels with significant group differences (*t*-test, EOS vs. typically developing [TD] controls; false discovery rate [FDR], *p_FDR_* < 0.05) were surrounded by black contours. Normative pediatric connectome was constructed using resting-state functional MRI data and diffusion tensor imaging data across TD controls. Connectivity degree of each parcel was computed by summing all edges of its binarized connectivity profiles. *Middle:* cortical parcels showing significant disease epicenters (Pearson’s correlation; spin permutation test, 10,000 times, *p_spin_* < 0.05) were surrounded by black contours. *Right:* the cortical parcels were then ranked and colored by their epicenter scores, i.e., correlation coefficients. Warm color refers to cortical thickening in patients, and cool color refers to cortical thinning.

### Functional and structural disease epicenters for EOS

Disease epicenters were located through evaluating whether the cortical thickness alterations in EOS patients were related to the normative pediatric network organization (**Figure 1**). Herein the normative pediatric network organization refers to the averaged functional connectome derived from resting-state functional MRI data across TD, as well as the averaged structural connectome derived from diffusion tensor imaging data in the same sample. Parcels with connectivity profiles significantly related to abnormal patterns of cortical thickness in patients were identified as EOS-specific epicenters. Significance was assessed by using spin permutation tests (10,000 times). Regions with positive values in disease epicenter maps refer to the connectivity profiles spatially resembling cortical thickening patterns in patients, while negative values resemble cortical thinning patterns. With respect to functional connectivity, disease epicenters of cortical thickness reduction were mainly located in the ventral attention network, frontoparietal control network and DMN, while epicenters of increased cortical thickness were in the visual network and sensorimotor network (*p_spin_* < 0.05). For structural connectivity, disease epicenters were similar to the functional epicenters (**Table S3**), but had less epicenters in the DMN and sensorimotor network and more epicenters in the visual network. The findings of EOS-specific epicenters held true when compared to replicated epicenters generated using normative adult connectivity data from the Human Connectome Project (**Figure S2**) (38).

To evaluate whether disease epicenters shifted with increasing age, we further conducted an explorative analysis of disease epicenter dynamics (See Supplement). Briefly, we found that the disease epicenter map gradually faded from childhood to adolescence (**Figure S6**). Additionally, to test whether disease epicenters in patients with EOS differed from adult-onset schizophrenia, we calculated the disease epicenters of adult-onset schizophrenia (**Figure S4**) using the Cohen’s d map for adult-onset schizophrenia from the ENIGMA Toolbox (37). We observed that the functional epicenter pattern in adult-onset schizophrenia was significantly correlated with functional epicenters in patients with EOS (*r* = 0.6; *p_spin_* = 0.002), unlike the structural epicenter pattern (*r* = 0.3; *p_spin_* = 0.1). Notably, patients with adult-onset schizophrenia only showed negative epicenters in association cortices involved in the DMN, frontoparietal control network, limbic network, and ventral attention network (*p_spin_*< 0.05) but not positive epicenters, indicating a reserved association end of cortical thickness reduction and vanished sensorimotor end of cortical thickness increase in adult-onset schizophrenia relative to patients with EOS.

### Neurodevelopment, functional systems, and cognitive embedding

To further embed disease epicenters within the neurodevelopmental, functional, and cognitive continuum, we ranked 400 parcels in ascending order based on the epicenter values. Close correlations (**Figure 2A**) were detected between functional (*r* = 0.74; *p_spin_* < 0.0001) and structural (*r* = 0.55; *p_spin_* < 0.0001) disease epicenter axes and the neurodevelopmental axis suggested by a previous work (22).

**Figure 2.**
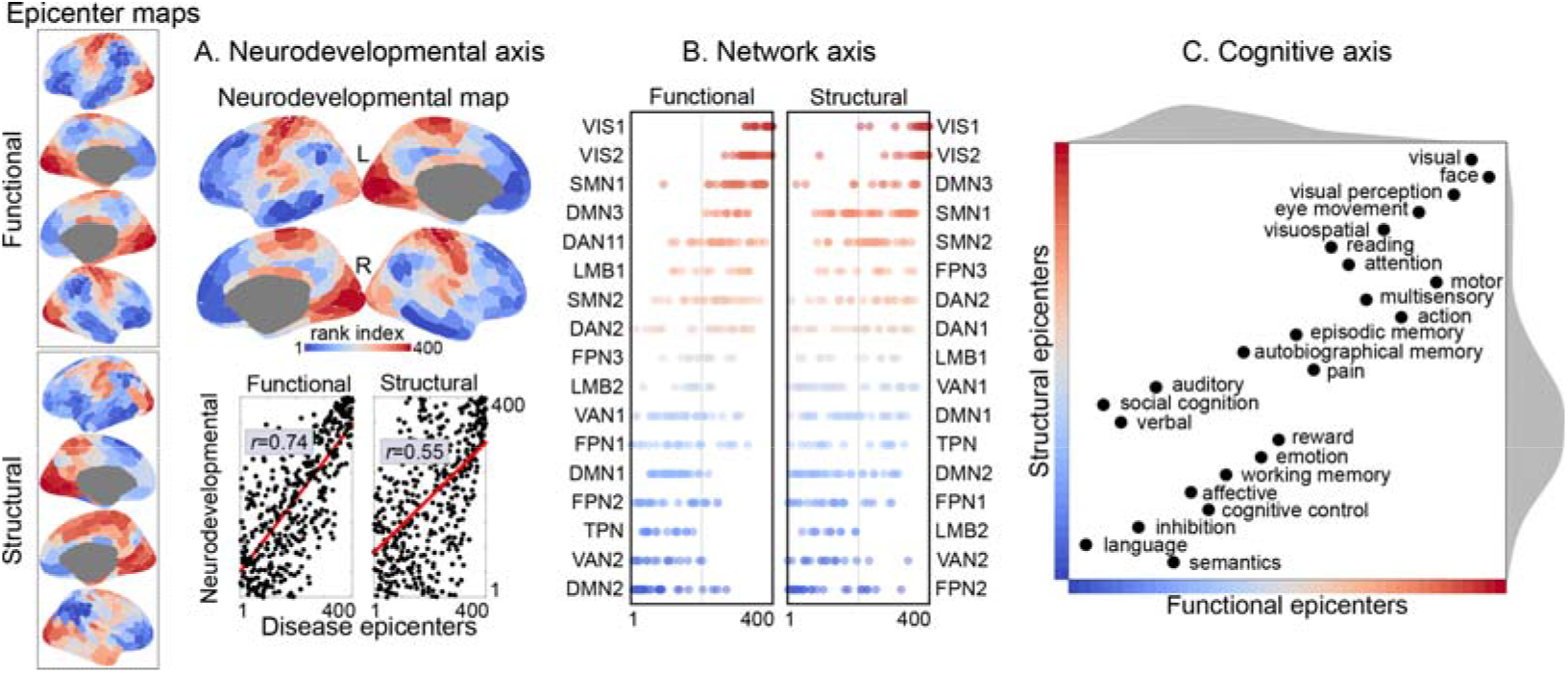
Associations with the neurodevelopmental, functional, and cognitive continuum. **(A)** Spatially correlating the neurodevelopmental axis with disease epicenter maps. The neurodevelopmental map was identified by a previous review (22) and was ranked by their developmental hierarchy. Pearson’s correlations were calculated between ranked disease epicenter map and ranked neurodevelopmental map (*p_spin_*< 0.05, 10,000 times). **(B)** Functional system distributions of disease epicenter maps. Ranked cortical parcels were subdivided into 17 networks according to a prior parcellation atlas (45), including the central visual network (VIS1), peripheral visual network (VIS2), sensorimotor A network (SMN1), sensorimotor B network (SMN2), dorsal attention A network (DAN1), dorsal attention B network (DAN2), ventral attention A network (VAN1), ventral attention B network (VAN2), orbitofrontal limbic network (LMB1), temporal-pole limbic network (LMB2), frontoparietal control A network (FPN1), frontoparietal control B network (FPN2), frontoparietal control C network (FPN3), default mode A network (DMN1), default mode B network (DMN2), default mode C network (DMN3), and temporoparietal network (TPN). **(C)** Cognitive term distributions in the epicenter space. In the two-dimensional epicenter space, 24 points refer to 24 cognitive terms. The location of each term was estimated by the association between its activation map and 40 functional (x-axis) and structural (y-axis) epicenter bins. The density map plotted by kernel density estimation function represents the distribution of *z*-statistic values for epicenter bins, capturing epicenter-cognition associations.

By assigning 400 areas to 1 of 17 functional systems (45), we found that visual and sensorimotor systems defined the positive end of the epicenter axis and frontoparietal control network and DMN defined the negative end (**Figure 2B**). The attention system was located in the middle of the axis. This functional system arrangement of the epicenter axis was aligned with hierarchical functional system development spanning from systems that process concrete and extrinsic information to systems subserving attention, then to systems linked to abstract and intrinsic processing (46).

Next, we identified cognitive implications of disease epicenters by conducting a meta-analysis on task-specific functional activations for 24 cognitive terms using the NeuroSynth database (47). In the two-dimensional space framed by functional (x-axis) and structural (y-axis) epicenters (**Figure 2C**), each cognitive term was situated by the association with disease epicenter bins assessed by z-statistics. A visual-memory-affective-language transition was observed for both axes. However, we found the social cognition term as an outlier of the correlation between structural and functional epicenters (*r_s_* = 0.89; *p* < 0.0001; **Figure S3**) by estimating the bootstrapped Mahalanobis distance (48), indicating different locations of the social cognition between structural and functional epicenter axes.

### Transcriptomic decoding

To further investigate the transcriptomic expression patterning underlying disease epicenters, we used high-resolution whole-brain microarray gene expression data derived from a postmortem brain transcriptomic dataset [AHBA; N = 6] (30) and transformed the data into a 400 × 15,631 (parcels × genes) matrix of normalized transcriptional levels via the abagen toolbox (41). Next, PLS regression (49) was used to reveal statistically significant latent variables relating the transcriptional matrix to disease epicenter patterns. As shown in **Figure 3A**, the first latent variable (PLS1) represents a covarying pattern of gene expression weights and disease epicenter weights (*r* = 0.50; *p_spin_* < 0.0001) that captured 98% covariance (*p_spin_* < 0.0001). The covarying pattern between gene expression and epicenters showed a sensorimotor (PLS+) to association (PLS−) transition across the cortex (**Figure 3B**).

**Figure 3.**
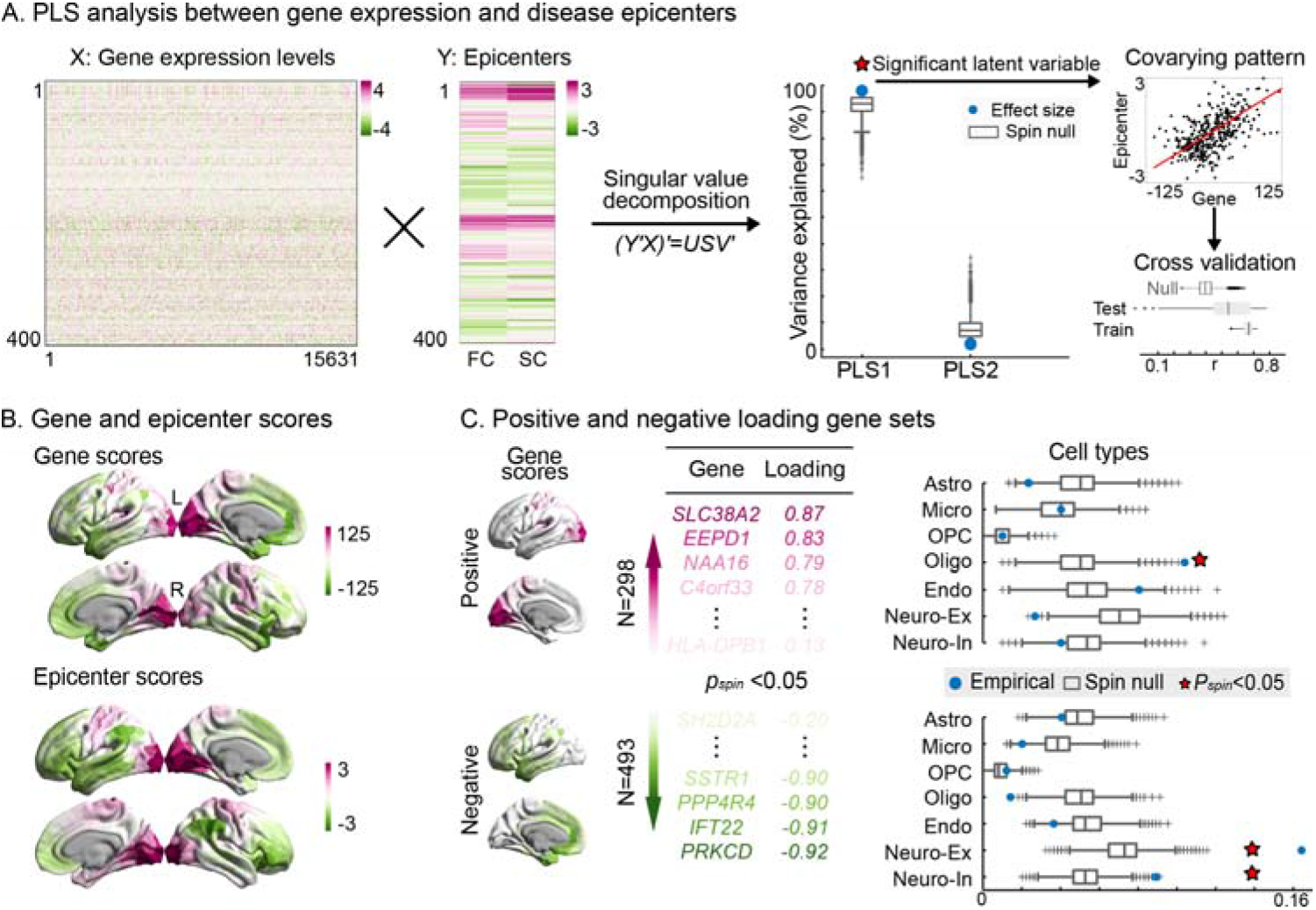
Underlying transcriptomic architecture. **(A)** Performing partial least squares (PLS) analysis to identify transcriptomic architecture underlying disease epicenters. Two input variables include a 400 × 15631 (parcels × genes) microarray gene expression matrix (30) and a 400 × 2 (functional and structural epicenters) epicenter matrix. After correlating the two variables across parcels and singular value decomposing (43), several latent variables capturing maximally covarying patterns within input variables were generated. Blue points refer to latent variables ordered by effect size of explained variance, and grey boxplots (the first, second (median) and third quartiles) refer to null distributions generated by spin permutations (10,000 times). The first latent (PLS1) that accounts for 98% of the covariance was statistically significant (*p_spin_* < 0.0001), capturing a covarying pattern of gene expression weights and disease epicenter weights (*r* = 0.50; *p_spin_* < 0.0001). The covarying pattern was cross-validated (100 times) by constructing the training set (Pearson’s correlation, mean ± SD; *r* = 0.66 ± 0.04) with 75% parcels closest to a randomly chosen parcel, and the testing set (*r* = 0.52 ± 0.19) with the remaining 25%. The significance was determined by spin permutations (*p_spin_* < 0.05, 10,000 times). **(B)** Covarying gene and epicenter score maps. Gene and epicenter scores were obtained by projecting input data onto gene and epicenter weights of PLS1. Warm color represents the PLS+ pattern, while cool color represents the PLS− pattern. The deeper color refers to the deeper extent of the parcel expresses the covarying pattern. **(C)** Gene loadings and involved cell types. Gene contributions were determined by gene loadings, which were calculated by projecting gene expression matrix onto gene score map of the PLS1. There were 298 PLS+ and 493 PLS− genes significantly contributing to the PLS1 (*p_spin_* < 0.05, 10,000 times). In the right panel, a blue point refers to the ratio of PLS+ (PLS−) gene set preferentially expressed in a certain cell type estimated by a cell-type deconvolution approach (32). The null distributions were constructed by randomly selection of all genes (*p_perm_* < 0.05, 10,000 times). Astro, astrocyte; Micro, microglia; OPC, oligodendrocyte precursor; Oligo, oligodendrocyte; Endo, endothelial; Neuro-ex, excitatory neurons; Neuro-in, inhibitory neurons.

To index the contribution of each gene to PLS1, we computed gene loadings by correlating the gene score map and gene expression matrix. We found a total of 791 genes, including 298 PLS+ and 493 PLS− genes (**Figure 3C**), that contributed significantly to the latent variable (*p_spin_* < 0.05). The PLS+ gene set represents that these genes were more expressed in positive epicenter regions, and vice versa. Next, we computed the ratio of these genes that are preferentially expressed in specific cell types, including astrocytes, microglia, oligodendrocyte precursors, oligodendrocytes, endothelial cells, excitatory neurons, and inhibitory neurons by using cell-specific aggregate gene sets derived from previous human postmortem single-cell and single-nucleus RNA sequencing studies (43). We observed that PLS+ genes had a significantly stronger expression of oligodendrocytes (*p_spin_* < 0.0001), while PLS− genes were more highly expressed in excitatory (*p_spin_* < 0.0001) and inhibitory neurons (*p_spin_* < 0.0001). To further identify the biological processes involved in these epicenter-associated gene sets, we aligned various enrichment terms, such as gene ontology biological processes, with PLS+ (PLS−) gene lists using the Metascape toolbox (44) (See Supplement).

### Associations with major brain disorders and HAR genes

To further link epicenter-related genes with major brain disorders, we intersected the PLS+/ PLS− gene lists (*p_spin_*< 0.05) and the genes differentially expressed in postmortem brain tissue measurements of mRNA (*p_FDR_* < 0.05) in five major neuropsychiatric disorders, including schizophrenia, autism spectrum disorder, bipolar disorder, major depressive disorder, and alcohol abuse disorder (34). Inflammatory bowel disease was also included as a non-neural control. Given the potential impact of gene outliers (50), we performed Spearman’s correlation analysis between PLS loadings and disease-specific differential gene expression (DGE) values. A positive DGE value of a gene indicates upregulation of transcriptomic expression in a disorder, while a negative DGE value indicates downregulation. We found that epicenter-related gene loadings were significantly correlated with DGE values of schizophrenia (*r_s_*(*_93_*) = 0.30; *p_perm_* = 0.003), autism spectrum disorder (*r_s_*(*_85_*) = 0.52; *p_perm_* < 0.0001), and bipolar disorder (*r_s_*(*_29_*) = 0.50; *p_perm_* = 0.005; **Figure 4A**), which indicated that PLS+ (PLS−) genes were linked with gene upregulation and downregulation, respectively, in these disorders. To further account for potential categorical aspects of PLS loadings, we binarized PLS loadings and DGE values according to the signs, then performed Kendall’s correlation analysis. We detected a convergingly positive correlation between binarized PLS loadings and binarized DGE values in schizophrenia (*r_k_*(*_93_*) = 0.45; *p_perm_* < 0.0001), autism spectrum disorder (*r_k_*(*_85_*) = 0.65; *p_perm_* < 0.0001), and bipolar disorder (*r_k_*(*_29_*) = 0.92; *p_perm_* < 0.0001).

**Figure 4.**
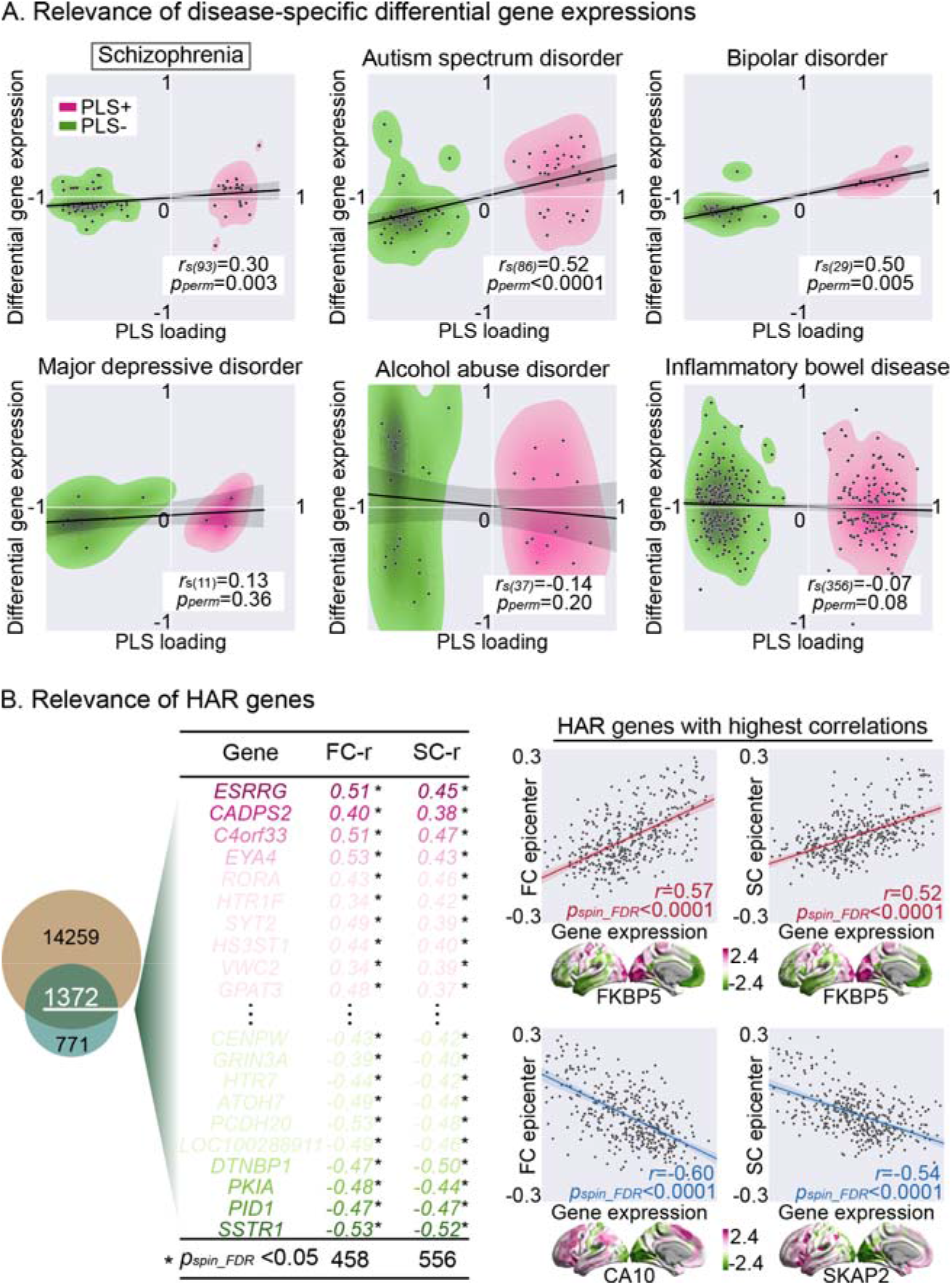
Relevance with major brain disorders and human accelerated region (HAR) genes. **(A)** Correlations between PLS loadings and histological measures of differential gene expressions of six brain disorders (34). The number of genes significantly related with epicenters (*p_spin_* < 0.05) and disorders (*p_FDR_* < 0.05) is 94 for schizophrenia, 86 for autism spectrum disorder, 30 for bipolar disorder, 12 for major depressive disorder, 38 for alcohol abuse disorder, and 357 for inflammatory bowel disease. Spearman’s correlation was calculated between PLS loadings and differential gene expression values across genes. The significance was estimated by permutation test (*p_perm_* < 0.05, 10,000 times). **(B)** Correlation between epicenter maps and gene expression maps of HAR genes. A total of 2143 HAR genes were selected (51) and overlapped with 15631 AHBA genes, resulting 1372 genes with transcription data. In the left panel, these genes were then ranked by PLS loadings. Pearson’s correlations (FC-r/ SC-r values) were then computed between functional/ structural epicenter maps and gene expression maps of these genes. The asterisk represents a significant association evaluated by spin permutation test (*p_spin_FDR_* < 0.05, 10,000 times), corrected by FDR method. The highest positive and negative correlations were shown in the right panel.

To determine the relationship between epicenter-related genes and HAR genes that dominate human-specific brain development implicated in schizophrenia, we selected 2143 HAR genes from a previously study (51) and overlapped the HAR genes with 15,631 AHBA genes, resulting in 1372 genes with transcription data (**Figure 4B**). Next, we ranked these genes using epicenter-related PLS loadings. We observed that 31 HAR genes were intersected with PLS+ genes (10% overlap) and 70 genes with PLS− genes (14% overlap). Gene expression maps involving 458 of 1372 HAR genes were significantly related to the functional epicenter map (*p_spin_* < 0.05 [FDR corrected]) and 556 HAR genes with structural epicenters (*p_spin_* < 0.05 [FDR corrected]). The highest positive correlations were detected in FKBP5 for both functional (*r* = 0.57; *p_spin_FDR_* <0.0001) and structural epicenters (*r* = 0.52; *p_spin_FDR_* <0.0001), while the highest negative correlations were detected in CA10 for functional epicenters (*r* = −0.60; *p_spin_FDR_* <0.0001) and SKAP2 for structural epicenters (*r* = −0.54; *p_spin_FDR_* <0.0001).

## Discussion

In this study, we investigated disease epicenters of cortical thickness alterations in patients with EOS and found epicenters of thickness reductions in association areas and epicenters of increased thickness in sensorimotor regions. The sensorimotor-to-association (S–A) spatial pattern aligns with the human neurodevelopmental axis from childhood to adolescence (22), and reflected a cognitive continuum from visual, motor, memory, affective to language. We observed a set of axon-related genes associated with the sensorimotor-end and synapse-related genes involved in the association end of our epicenter map using transcriptomic decoding. These epicenter-related gene weights were related to dysregulated gene expression of schizophrenia, autism spectrum disorder, and bipolar disorder, but not depression, substance use disorder, and inflammatory bowel syndrome. Moreover, our epicenter map was closely linked with the transcriptomic architecture of HAR genes that may harbor common genetic determinants within neurodevelopmental diseases. These results suggest a S–A spatial axis of disease epicenters that differentiates cortical alterations of sensorimotor and association regions to two poles and illustrate distinct microscale transcriptomic architectures underlying cortical thinning and thickening processing in EOS.

Based on the assumption that pathologic changes spread between anatomically or functionally connected brain regions (52), we used the epicenter model to identify probable pathologic origins in EOS. Disease epicenters refer to brain regions influenced earlier by schizophrenia, and could serve as a gateway affecting downstream hub nodes via their connections (7). Given both cortical increases and decreases in EOS, we measured positive and negative disease epicenters for the first time, rather than just one epicenter end of cortical decreases (7). Particularly, if the brain connectivity profile of a region is highly negatively correlated with a cortical abnormality map of schizophrenia (negative epicenters), this region is strongly connected to other high-thinning regions and weakly connected to other low-thinning regions. Conversely, a positive correlation (positive epicenters) represents that this region is strongly (weakly) connected to other high-thickening (low-thickening) regions. We found EOS epicenters of cortical thinning in association cortices, including the orbitofrontal cortex, inferior parietal lobule, and temporal lobe, while disease epicenters of cortical thickening were found in sensorimotor cortices, including visual and sensorimotor-related regions. The association epicenters of cortical thinning support previous findings of frontotemporal epicenters of grey matter volume loss (6, 8) and cortical thinning (7) in adult-onset schizophrenia. Generally, patients with schizophrenia exhibited widespread and progressive cortical thinning relative to healthy controls (53), with the largest effect sizes for the frontotemporal cortex (54).

Conversely, increased cortical thickness in TD children appears to reflect higher polygenic risk for schizophrenia (55). Excessive thickening of the cortex has been found in healthy individuals with increased schizotypy scores which is thought to be an abnormal neurodevelopmental trait (56). Accordingly, this unexpected cortical thickening pattern might serve as EOS-specific developmental abnormality relative to adult-onset schizophrenia. Alternatively, a few studies have suggested that patients with an early stage of schizophrenia have cortical thickening in the parietal lobule and occipital pole (57). Consequently, albeit not confirmed by longitudinal data, this thickening pattern might be a precursor of the disease that gradually disappears with disease progression. Overall, the current findings reveal convergent cortical thickness reductions of association cortices in EOS, and further underscore thickening of sensorimotor cortices in the early stage of schizophrenia.

Orbitofrontal epicenters of cortical thinning were found in functional epicenters but not in structural epicenters. These regional differences between functional and structural epicenters were linked to social cognition functions as probed by a meta-analysis on task-specific functional activations. The differences between functional and structural epicenters may result from different neurobiological and functional mechanisms of underlying connectome construction. First, the functional connectome characterizes an indirect relationship between regions relative to structural connectome. For example, an indirect relationship could be produced by polysynaptic connections, thus reflecting inherent characteristics of the functional connectome (58). Indeed, although most studies had consistent findings between functional and structural epicenters (6, 7, 17), a recent longitudinal study found that pathology of psychotic illness spreads through structural connectivity, but not functional connectivity. Thus, our orbitofrontal finding of functional epicenters without structural foundation should be interpreted with care. In addition, the absence of inter-hemisphere structural connections could also lead to potential loss of anatomic and spreading relevant information. From a methodologic standpoint, the reduced number of structural epicenters might be caused by calculating whole-cortex epicenters using intra-hemispheric structural connectome. Another explanation for the difference between functional and structural epicenters is structure-function uncoupling of the transmodal association cortex (59). Previous studies have reported structure-function coupling of the primary unimodal cortex and structure-function uncoupling of the transmodal association cortex, and suggested that the different relation between structure and function in association areas may support flexible cognition and behavior in humans (59, 60). Consequently, it may be that rather than direct connections, more indirect functional associations between impacted regions are impaired, linked to downstream functional alterations in the social cognitive domain. Nevertheless, the difference between structural and functional epicenters can be attributed to multiple reasons and need further investigations.

As we hypothesized, the disease epicenter pattern in EOS aligns well with the human neurodevelopmental hierarchy from childhood to adolescence (22), i.e., the S–A axis. Sensorimotor regions are situated on one end and association regions on another. After cortical thickness reaches a peak in early childhood (61, 62), cortical thickness undergoes a protracted developmental decline from childhood to adolescence. Primary sensorimotor regions, including occipital, pre- and post-central, and medial temporal cortices undergo a rapid thinning in childhood (63), while exhibit minimal thinning in adolescence (64). Transmodal association cortices, including temporal, parietal, and frontal cortices, exhibit mild thinning or even localized thickness increases in childhood (65, 66) but are enhanced thinning during adolescence (67). Indeed, in our explorative analyses testing whether epicenters show patterns of disease progression as a function of age of onset, we noted that disease epicenters of EOS shifted and gradually faded out with increasing age. Specifically, disease epicenters of cortical thinning were predominantly in the frontotemporal regions in childhood, while the temporal epicenters shifted up in early adolescence. Sensorimotor and visual epicenters of cortical thickening disappeared in early adolescence. The S–A disease epicenter pattern was disorganized and spread to the entire cortex in late adolescence. This age-related divergent pattern of the epicenter is coincident with previous findings of unique epicenters for first-episode and early stages relative to chronic stages of schizophrenia (7). Disease epicenters of psychotic illness have been reported to longitudinally evolve with illness progression and antipsychotic exposure (8). Beyond the influence of illness and medication, our findings additionally reveal a potential neurodevelopmental effect on disease epicenters. A hypothesis worthy of evaluation is whether EOS patients show a delayed maturation of grey matter in sensorimotor cortices resulting in reduced cortical thinning in childhood and faded patterns in adolescence. Conversely, patients seem to have excessive maturation of grey matter in association cortices due to increased cortical thinning during childhood and adolescence. Taken together, our findings may yield new insight into cortical structural abnormalities in schizophrenia from a disturbed S–A developmental hierarchy and may motivate further work into the lifespan trajectories of schizophrenia.

Benefitting from imaging transcriptomics advancements and open resources (30, 31), we could investigate the potential microscopic neurobiological substrate underpinning the S–A epicenter pattern. We identified cortical expression of a weighted combination of genes that most collocated with the EOS epicenter pattern. Disease epicenters of cortical thickening were enriched for genetic signaling of oligodendrocytes, a type of non-neuronal cells involved in glial function, especially myelin production (68). In contrast, disease epicenters of cortical thinning colocalized with cortical expression of genes related to excitatory and inhibitory neurons rather than support cells. Indeed, neurodevelopmental plasticity during childhood and adolescence is associated with cellular and circuit refinement processes of glial, excitatory, and inhibitory cells (22). Oligodendrocytes induce myelination of sensorimotor regions in childhood (69), which could suppress synaptic plasticity to increase stability. Thus, excessive expression of oligodendrocytes-related genes appears to underpin the developmental delay of the sensorimotor pole in schizophrenia during childhood as we assumed (see above). Excitatory neurons (such as pyramidal neurons) are pruned and inhibitory neurons grows (such as parvalbumin interneurons) during brain maturation, resulting a decline of the cortical excitatory/inhibitory ratio (70). This decline increases microcircuit signal-to-noise ratio and shifts the balance of circuit activity from spontaneous to evoked, served as a hallmark of neurodevelopmental plasticity (71). During the neurodevelopmental critical period, excitatory/inhibitory abnormalities of association regions have been suggested to underlie the emergence of psychopathology (70, 72, 73). Accordingly, pathologic abnormality in association regions in schizophrenia is likely to be attributed to dysregulated expressions of excitatory and inhibitory neurons-related genes. These findings provide micro-level evidence for a developmental component involved in pathologic origins of schizophrenia. Future work should be conducted across scales to further link microscale cellular and molecular changes with macroscale cortical abnormalities in schizophrenia.

According to the neurodevelopmental hypothesis of schizophrenia (3), early neurodevelopmental disturbances contribute to the pathogenesis of schizophrenia. We indeed observed that the neurodevelopmentally rooted S–A epicenter axis was associated with genes differentially expressed in postmortem case-control studies of schizophrenia. Here cortical thickening in schizophrenia was related to disease-specific gene upregulation and cortical thinning to downregulation. However, the epicenter axis was also associated with dysregulated genes in autism spectrum and bipolar disorders. Coincidently, an emerging view proposed that neurodevelopmental-related mental diseases including schizophrenia could be conceptualized as lying on an etiologic continuum (25). To be specific, most symptom domains, including cognitive impairment, negative symptoms, and positive symptoms, are shared among these psychiatric diseases. Although cognitive impairments are most severe and pervasive in autism spectrum disorder, negative symptoms are more pronounced in schizophrenia. Positive symptoms are more apparent in bipolar disorder (34). These subtle differences essentially reflect the degree and timing of abnormal neurodevelopment. This notion of a neurodevelopmental continuum has been supported by transdiagnostic genetic loci that are implicated in synaptic development and plasticity (26, 74). Shared transcriptional dysregulation further indicates polygenic overlap across these disorders, which converges in common neurobiological pathways (34). The current findings provide new evidence for the neurodevelopmental continuum, and indicate that the S–A epicenter axis might serve as a converging pathophysiologic framework across schizophrenia, autism spectrum disorder, and bipolar disorder.

Recently, the neurobiological continuum across psychiatric disorders was explained by an evolutionary hypothesis, suggesting these mental illnesses emerge as costly by-products of human evolution (27). With the neuroscientific and technological advances, genes located in HARs of the genome can be identified and regarded as evolutionary markers, e.g., it can be investigated whether they are engaged in human-specific neurodevelopment and outcomes (75). These HAR genes have been suggested to have a vital role in human brain development (76) and may induce multiple brain disorders (77). As a previous review suggested (29), HARs are involved in the genetic signature of neurodevelopmental-related psychiatric disorders including schizophrenia. Accordantly, we found an association between the S–A epicenter axis of EOS and brain expression maps of HAR genes. This finding links macroscale pathologic phenotypes of schizophrenia with microscale transcriptomics of HAR genes, partly supporting the evolution hypothesis. Yet, the neurodevelopmentally rooted epicenter axis embeded schizophrenia, autism spectrum disorder, and bipolar disorder along a common continuum, and may shed new insight into bridging neurodevelopment, schizophrenia, and evolution. Of course, further studies on non-human primate datasets are highly recommended to explore the relationship between human evolution and the neurodevelopmental disease model.

Our study had several technological limitations that need to be considered. First, present findings of epicenters are based on correlational analyses in cross-sectional data, making it impossible to infer the causality of cortical thickness alterations. We cannot resolve whether there is another underlying mechanism potentiating brain deformation beyond functional and structural connectivity. Although age-related subgroup comparisons suggested a dynamic wave of disease epicenters, the relatively small sample size for these subgroups were underpowered to uncover stable and robust findings. Moreover, the cross-sectional design could be confounded by multiple aspects of individual variability. Next, transcriptome-neuroimaging associations were established on prior adult gene expression data without psychiatric diagnoses (30), hindering our examination of the relationships across groups. Even though we observed the associations between gene weights and dysregulated gene expression of the postmortem samples from patients with schizophrenia (34), these correlational analyses still cannot provide direct evidence for our findings. Third, restricted clinical variables were collected in this study, that might account for the lack of a relationship between clinical behaviors and brain abnormalities. Further studies are recommended to explore the relationship between disease epicenter patterns and clinical behaviors of patients. Fourth, the set of HAR genes was selected according to a previous study (51), while there are some other alternative approaches (78). Although these different selections of HAR genes result consistent findings as a previous study suggested (35), future works should include a more comprehensive set of HAR genes. Last, we only involved EOS patients to study neurodevelopmental disease, but not other disorders, such as autism spectrum disorder. Despite the observed microscale association with autism spectrum and bipolar disorders, future studies are needed to directly test the generalizability of our epicenter model in diverse neurodevelopmental diseases.

In conclusion, our study revealed a sensorimotor-to-association disease epicenter map that differentiated cortical thickness alterations in EOS and uncovered the underlying microscale processes through transcriptomic analyses. Our findings suggest developmentally rooted pathologic origins of schizophrenia during brain maturation. Broadly, this study provides a unified framework to understand the etiology of schizophrenia and other neurodevelopmental-related psychiatric disorders. The framework may help identify neurobiological markers critical for early diagnosis and intervention.

## Data Availability

Functional and structural disease epicenter maps of EOS and other data supporting our findings are available at https://github.com/Yun-Shuang/Neurodevelopmentally-rooted-epicenters-in-schizophr <u>enia</u>. Human gene expression data are available at https://human.brain-map.org/. Additional information can be made available upon reasonable request to the authors.

## Code Availability

Custom code was made publicly available under https://github.com/Yun-Shuang/Neurodevelopmentally-rooted-epicenters-in-schizophr enia. Epicenter calculation was based on ENIGMA Toolbox (https://enigma-toolbox.readthedocs.io/en/latest/); structural covariance gradients calculation was based on BrainSpace (https://brainspace.readthedocs.io/en/latest/); cognitive meta-analysis code was adapted from https://github.com/CNG-LAB/cngopen/blob/main/transdiagnostic_gradients/Scripts/H ettwer2022_Figure2_Transdiagnostic_Gradients.m; gene expression analyses were performed by the abagen toolbox (https://abagen.readthedocs.io/), combined with code under https://github.com/netneurolab/hansen_genescognition; gene enrichment analyses by metascape (https://metascape.org/gp/index.html#/main/step1); statistically analyses were carried out by BrainStat (https://github.com/MICA-MNI/BrainStat); visualizations were based on workbench (https://www.humanconnectome.org/software/connectome-workbench) and ggseg (https://ggseg.github.io/ggseg/), combined with ColorBrewer (https://github.com/scottclowe/cbrewer2).

## Acknowledgements

We are grateful to all the participants and their guardians in this study. We thank International Science Editing (http://www.internationalscienceediting.com) for editing this manuscript. This work was supported by STI 2030—Major Projects 2022ZD0208900, the National Natural Science Foundation of China (62333003, 62036003, 82121003, 62373079, 62073058), Medical-Engineering Cooperation Funds from University of Electronic Science and Technology of China (ZYGX2021YGLH201). Y-S.F. was also funded by the China Postdoctoral Science Foundation (2023M740524) and Sichuan Province Innovative Talent Funding Project for Postdoctoral Fellows. S.L.V. was also funded in part by Helmholtz Association’s Initiative and Networking Fund under the Helmholtz International Lab grant agreement InterLabs-0015, and the Canada First Research Excellence Fund (CFREF Competition 2, 2015-2016) awarded to the Healthy Brains, Healthy Lives initiative at McGill University, through the Helmholtz International BigBrain Analytics and Learning Laboratory (HIBALL). M.D.H. was funded by the Max Planck Society and the German Ministry of Education and Research (BMBF).

