## Supplement for "Neurodevelopmentally rooted epicenters in schizophrenia: sensorimotor-association spatial axis of cortical thickness alterations"

Running title: Brain epicenters of early-onset schizophrenia

**Yun-Shuang Fan** ^1,2^ PhD, **Yong Xu** ^3^ PhD, **Meike Dorothee Hettwer** ^2,5-7^ PhD, **Pengfei Yang** ^1^ MD, **Wei Sheng** ^1^ PhD, **Chong Wang** ^1^ PhD, **Mi Yang** ^1^ PhD, **Matthias Kirschner** ^8^ PhD, **Sofie Louise Valk** ^2,5*#^ PhD, **Huafu Chen** ^1,4*#^ PhD

*1.The Clinical Hospital of Chengdu Brain Science Institute, School of Life Science and Technology, University of Electronic Science and Technology of China, Chengdu, China; 2. Otto Hahn Group Cognitive Neurogenetics,* *Max Planck Institute for Human Cognitive and Brain Sciences, Leipzig, Germany; 3. Department of Psychiatry, First Hospital/First Clinical Medical College of Shanxi Medical University, Taiyuan, China; 4. MOE Key Lab for Neuroinformation, High-Field Magnetic Resonance Brain Imaging Key Laboratory of Sichuan Province,* *University of Electronic Science and Technology of China, Chengdu, China; 5. Institute of Neuroscience and Medicine (INM-7: Brain and Behavior), Research Centre Jülich, Jülich, Germany; 6. Max Planck School of Cognition, Leipzig, Germany; 7. Institute of Systems Neuroscience, Medical Faculty, Heinrich Heine University Düsseldorf, Düsseldorf, Germany; 8. Division of Adult Psychiatry, Department of Psychiatry, University Hospitals of Geneva, Geneva, Switzerland.*

* Both last co-authors contributed equally.

\# Corresponding authors:

Huafu Chen, & Sofie Louise Valk,.

**Supplement**

**Methods**

**Participants**

A total of 199 pediatric participants aged 7 to 17 years were recruited from the First Hospital of Shanxi Medical University, including 99 drug-naïve first-episode early-onset schizophrenia (EOS) patients and 100 typically developing (TD) controls. Patients were diagnosed by at least one senior psychiatrist (Y.X.). The severity of clinical symptoms was assessed using the positive and negative syndrome scale (PANSS) (N=71). Exclusion criteria for patients included a history of neurological MRI anomalies, substance abuse, or illness duration > 1 year, the use of antipsychotic medication. Exclusion criteria for controls additionally included history or family history of psychiatric disorder. This retrospective study was approved by the Ethics Committee of the First Hospital of Shanxi Medical University. Informed consent was obtained from all individuals and their parents or legal guardians. From this original sample, four patients were excluded due to incomplete scanning data, and one patient was excluded due to poor quality of cortical parcellation. A final sample including 95 EOS patients and 99 demographically-matched TD controls were further analyzed in the study (See **Table S1** for detailed demographic data).

**Imaging data acquisition**

Multimodal imaging data were acquired on a 3 Tesla Siemens MAGNETOM Verio scanner at the First Hospital of Shanxi Medical University. T1-weighted data were acquired via a three-dimensional fast spoiled gradient-echo sequence [repetition time (TR) = 2,300 ms; echo time (TE) = 2.95 ms; flip angle = 9°; matrix = 256 × 240; slice thickness = 1.2 mm (no gap); and voxel size = 0.9375 × 0.9375 × 1.2 mm^3^, with 160 axial slices]. Resting-state fMRI (rs-fMRI) data were obtained using a two-dimensional echo-planar imaging sequence [TR = 2,500 ms; TE = 30 ms; flip angle = 90°; matrix = 64 × 64; number of volumes = 198; slice thickness = 3 mm (1 mm gap); and voxel size = 3.75 × 3.75 × 4 mm^3^, with 32 axial slices]. Diffusion tensor imaging (DTI) scans were acquired using an echo-planar pulse sequence with the following parameters [TR = 6,000 ms; TE = 90 ms; flip angle = 90°; matrix = 128 × 128; number of volumes = 39; slice thickness = 3 mm (3 mm gap); and voxel size = 1.875 × 1.875 × 3 mm^3^, with 45 axial slices; b = 1000s/mm^2^ for 36 diffusion directions, and three b = 0 s/mm^2^ images].

**Cortical thickness estimation**

All T1-weighted data were preprocessed with FreeSurfer package (v7.1.0, <http://surfer.nmr.mgh.harvard.edu/>), including cortical segmentation and surface reconstruction. Vertex-level cortical thickness values were estimated using the distance between the white and the pial surfaces. Next, surface meshes (∼80,000 vertices per mesh) were down sampled into a 400-parcel cortical ‘Schaefer’ parcellation atlas by averaging vertex-wise cortical thickness values within each parcel. For each parcel, cortical thickness features were compared between the EOS and TD groups using two-sample *t*-tests (sex and age were regressed out as covariates), corrected for multiple comparisons by the false discovery rate (FDR) procedure . To account for the nonlinear pattern of cortical development , we conducted a validation analysis of group comparison by using two-sample *t*-tests with sex, age and age squared regressed out as covariates.

**Normative pediatric connectivity matrix construction**

rs-fMRI data were preprocessed with the CBIG pipeline (https://github.com/ThomasYeoLab/CBIG) based on FSL (v5.0.9) and FreeSurfer (v7.1.0), which included removal of the first four volumes, slice-timing, motion correction, boundary-based registration to structural images, covariates regression, and bandpass filtering (0.01–0.08 Hz). These functional data were then down sampled into the same 400-parcel cortical ‘Schaefer’ parcellation atlas as T1-weighted data. Pearson's correlations were computed between each pair of time series of 400 parcels to generate a 400 × 400 connectivity matrix for each subject. Negative connections of the matrix were set to zero as previously suggested . Next, the matrix was then *z*-transformed and aggregated across all TD controls to construct a group-average functional connectome, i.e., normative functional connectivity matrix.

DTI data were preprocessed with FSL (FMRIB Software Library v5.0.9, http://www.fmrib.ox.ac.uk/fsl) and the diffusion toolkit, including eddy current correction, diffusion tensor model estimation and whole-brain fiber tracking. In particular, the tractogram was generated with 40 million streamlines, with a maximum tract length of 250 and a fractional anisotropy cutoff of 0.06 as previously suggested . Spherical-deconvolution informed filtering of tractograms was then used to reconstruct whole-brain streamlines weighted by the cross-sectional multipliers. Reconstructed streamlines were mapped onto 400 parcels to create structural connectivity matrices, where connection was defined by the number of streamlines between two regions (i.e., fiber density). Normative structural connectivity connectome was created by aggregating the connectivity matrices across all TD controls, and was log transformed to reduce connectivity strength variance.

**Statistical null models establishing**

The statistical significance of spatial correlations involved in following analyses was assessed using spin permutation tests that account for spatial autocorrelation . Null models were created by projecting the center spatial coordinates of cortical parcels onto the surface spheres via randomly sampled rotations. These null distributions of correlation coefficients that were generated by spatially permuted cortical maps were then compared with the empirical correlation coefficients, resulting the statistical significance, *p_spin_* value.

**Neurodevelopment, functional systems, and cognitive embedding**

To delineate the relevance between disease epicenters and neurodevelopment, functional systems, and cognitive functions, we ranked all 400 parcels in ascending order based on their correlation coefficients, refer to degrees of epicenter. First, the neurodevelopmental axis was aligned and compared with functional and structural epicenters by using spatial correlations (*p_spin_* < 0.05, 10,000 times). The ranked 400 parcels were then assigned to one of 17 functional systems including the central visual network (VIS1), peripheral visual network (VIS2), sensorimotor A network (SMN1), sensorimotor B network (SMN2), dorsal attention A network (DAN1), dorsal attention B network (DAN2), ventral attention A network (VAN1), ventral attention B network (VAN2), orbitofrontal limbic network (LMB1), temporal-pole limbic network (LMB2), frontoparietal control A network (FPN1), frontoparietal control B network (FPN2), frontoparietal control C network (FPN3), default mode A network (DMN1), default mode B network (DMN2), default mode C network (DMN3), and temporoparietal network (TPN). Next, we discretized ranked parcels into 40 equally sized bins and combined parcels in the same bin to a joint brain map. We conducted a meta-analysis on task-specific brain activations for 24 cognitive terms using the NeuroSynth database . The associations between joint maps and 24 cognitive-related maps were assessed by *z*-statistics. For visualization, each cognitive term was located into the two-dimensional epicenter space according to their z values with functional (x-axis) and structural (y-axis) epicenter maps.

**Gene expression data preprocessing**

Regional microarray expression data were obtained from 6 post-mortem brains (1 female, age = 42.50 ± 13.38) provided by the Allen Human Brain Atlas (AHBA, [https://human.brain-map.org](https://human.brain-map.org/); ). Data were processed with the abagen toolbox (version 0.1.3; <https://github.com/rmarkello/abagen>; ) using a 400-region volumetric atlas in MNI space . First, microarray probes were reannotated using data provided by a previous work ; probes not matched to a valid Entrez ID were discarded. Next, probes were filtered based on their expression intensity relative to background noise , such that probes with intensity less than the background in >=50.00% of samples across donors were discarded. When multiple probes indexed the expression of the same gene, we selected and used the probe with the most consistent pattern of regional variation across donors (i.e., differential stability; ). The MNI coordinates of tissue samples were updated to those generated via non-linear registration using the Advanced Normalization Tools (ANTs; <https://github.com/chrisfilo/alleninf>). Samples were assigned to brain regions in the provided atlas if their MNI coordinates were within 2 mm of a given parcel. To reduce the potential for misassignment, sample-to-region matching was constrained by hemisphere and gross structural divisions (i.e., cortex, subcortex/brainstem, and cerebellum, such that e.g., a sample in the left cortex could only be assigned to an atlas parcel in the left cortex; ). All tissue samples not assigned to a brain region in the provided atlas were discarded. Inter-subject variation was addressed by normalizing tissue sample expression values across genes using a robust sigmoid function . Normalized expression values were then rescaled to the unit interval. Gene expression values were then normalized across tissue samples using an identical procedure. Samples assigned to the same brain region were averaged separately for each donor and then across donors, yielding a 400 × 15631 matrix (regions × genes) of transcriptional levels.

**Transcriptomic genetic decoding**

We used partial least squares (PLS) analysis to decompose associations between gene expression (X_400 × 15631_) and epicenter maps (Y_400 × 2_) into orthogonal sets of latent variables with maximum covariance. Specifically, the column-wise *z*-scored gene expression matrix (X) was multiplied by the transposed epicenter matrix (Y), and then singular value decomposed as below:

(Y′X)′ = USV′

In the diagonal matrix S_2 × 2_, the i_th_ singular value represents the covariance between two singular vectors, i.e., U_15631 × 2_ (gene weights) and V_2 × 2_ (epicenter weights) that captured by the ith latent variable, where positively (negatively) weighted genes covary with positively (negatively) weighted epicenters. To evaluate the significance of the i_th_ latent variable (*p_spin_* < 0.05, 10,000 times), we estimated effect size as the ratio of squared i_th_ singular value to the sum of all squared singular values. The robustness of the PLS model (*p_spin_* < 0.05, 10,000 times) was evaluated by cross-validating (repeated 100 times) the correlation between gene and epicenter scores, in which the training set was constructed by 75% of brain regions closest in Euclidean distance to a randomly region and the testing set by the remaining 25% . Gene and epicenter scores were then computed as below:

Gene scores: M_400 × 2_ = X_400 × 15631_ U_15631 × 2_

Term scores: N_400 × 2_ =	Y_400 × 2_ V_2 × 2_

Here positively scored regions demonstrate the covariance between positively weighted genes and epicenters, and vice versa. For the latent variable i, gene loadings (15631 × 1) were calculated by Pearson's correlating gene scores M(i)_400 × 1_ with gene expression matrix X_400 × 15631_. Genes with high absolute loadings reflect high variance with the latent variable, indicating large contribution to the covarying pattern. The significances of gene loadings were evaluated by spin permutation tests (*p_spin_* < 0.05, 10,000 times).

**Identification of involving cell types and biological pathways**

To determine cell types involved in the gene sets identified by PLS analysis, we performed cell type deconvolution using cell-specific aggregate gene sets proposed by previous studies . Particularly, seven cell classes including astrocytes, endothelial cells, microglia, excitatory neurons, inhibitory neurons, oligodendrocytes, and oligodendrocyte precursors were estimated by using hierarchical clustering of regional topographies across all study-specific cell types in the AHBA (*p_spin_* < 0.05, 10,000 times), corrected for multiple comparisons (seven times) by the FDR procedure. To further determine biological pathways involved in the gene sets, we performed enrichment analysis using Metascape, a web-based portal that provides automated meta-analysis tools leveraging over 40 independent knowledgebases ([https://metascape.org/gp/index.html#/main/step1](https://metascape.org/gp/index.html" \l "/main/step1)) . We input PLS+ and PLS− gene sets into the Metascape website separately and thresholded resulting enrichment pathways (p < 0.05, FDR correction), then discarded discrete enrichment clusters.

**Association with differential gene expressions of major brain disorders**

To evaluate the relationship between epicenter-related loadings of the gene sets identified by PLS analysis and disease-specific gene dysregulation, we used postmortem brain tissue measurements of mRNA (*p_FDR_* < 0.05) in six brain disorders, including schizophrenia, autism spectrum disorder, bipolar disorder, major depressive disorder, alcohol abuse disorder (alcoholism), and inflammatory bowel disease . For example, we intersected PLS+/ PLS− gene lists and those genes that significantly differentially expressed in schizophrenia (1841 genes, *p_FDR_* < 0.05), then correlating their differential gene expression values with PLS loadings. Herein we performed Spearman’s correlation analyses to account for outlier genes , and the significance was evaluated by permutation tests (*p_perm_* < 0.05, 10,000 times). We also applied the same analyses on autism spectrum disorder-specific 2123 genes, bipolar disorder-specific 494 genes, major depressive disorder-specific 154 genes, alcoholism-specific 706 genes, and inflammatory bowel disease-specific 7201 genes (*p_FDR_* < 0.05) .

**Association with transcriptomic architecture of human accelerated regions (HARs)**

To test whether transcriptomic architecture of genes located in HARs of the genome underpin disease epicenter map, we used 2143 HAR genes defined by a previous study . In particular, these genes were selected out from 2737 HAR genes by using a comparative genome analysis . We filtered out 1372 HAR genes that were included in the AHBA dataset, and ranked them by epicenter-related PLS loadings. The overlap ratio between these genes and the gene sets identified by PLS analysis was calculated. Next, we used Pearson’s correlation analyses to examine the associations between brain expression maps of HAR genes and disease epicenter maps (*p_spin_* < 0.05, 10,000 times).

**Age-related disease epicenter mapping**

Having established the overall disease epicenter pattern for 7–17 years old patients with early-onset schizophrenia (EOS), we tested whether disease epicenters would shift with increasing age. First, we estimated individual-level cortical thickness deviations from typically developing (TD) controls by using w-score maps. In particular, parcel-wise w-score map for patient j was generated by the formula:

w-score(j) = (actual(j) − expected)/ SD

In this formula, the actual is patient’s cortical thickness, the expected is mean cortical thickness across TD, and the SD is the standard deviation of TD . Individualized epicenter map for each patient was calculated by spatially correlating w-score map with normative pediatric connectome and were then assigned to 17 functional systems. We divided all participants into three age-stage groups included childhood group (N = 69, 28 men, age range = 7–13 years), early adolescent group (N = 66, 21 men, age range = 14–15 years), and late adolescent group (N = 59, 25 men, age range = 16–17 years). See **Table S1** for detailed demographic information. Stage-specific disease epicenter maps were obtained by averaging individualized epicenter maps belonging to a subgroup.

Next, we also generated stage-specific disease maps by group-level epicenter mapping method. For each age stage, cortical thickness alterations in EOS were evaluated by performing two-sample *t*-tests between patients and controls with age and sex as covariates (false discovery rate [FDR]; *p_FDR_* < 0.05). Resulting *t* maps were then spatially correlated with stage-matched normative functional and structural connectomes to generate stage-specific disease epicenters (*p_spin_* < 0.05, 10,000 times).

**Results**

**Underlying biological processes**

To identify the biological processes involved in these epicenter-associated gene sets, we aligned various enrichment terms, such as gene ontology biological processes, with PLS+ (PLS−) gene lists using the Metascape toolbox . After computing accumulative hypergeometric *p*-values of enrichment terms (*p_FDR_* < 0.05) and discarding discrete clusters (**Figure S5**), we showed that PLS1+ genes were mainly enriched for gene ontology biological processes, such as chordate embryonic development, brain development, epithelia cell migration, tube morphogenesis, ear morphogenesis, centrosome cycle, and positive regulation of glial cell proliferation. The PLS1− genes were enriched for multiple pathways, including modulation of chemical synaptic transmission, transport along microtubule, regulation of membrane potential, neuronal system, adaptive immune system, nervous system development, potassium channels, antigen processing, intraflagellar transport proteins binding to dynein, and cholinergic synapse.

**Disease epicenters shifted from childhood to adolescence**

To map age-specific disease epicenters, we first calculated individualized epicenter map by correlating cortical thickness deviations of each patient with normative pediatric functional and structural connectomes. After assigning 400 parcels to one of 17 functional systems, network-level functional epicenter map was generated for each patient (**Figure S6A**). Ranking these networks by the overall disease epicenter axis, we found that the disease epicenter pattern gradually faded out with increasing age. Network-level structural epicenters were shown in **Figure S7**. Next, we divided all subjects into three groups (**Table S1**), including childhood group (7–13 years, N = 69), early adolescence group (14–15 years, N = 66), and late adolescence group (16–17 years, N = 59). By averaging network-level individualized epicenters within each group, we found that network location along the epicenter axis changed across different stages.

We also conducted group-level epicenter mapping analyses for each group to generate functional and structural disease epicenters within specific age range (**Figure S6B**). Negative disease epicenters were located in the default mode network and limbic network during childhood, while were superiorly shifted to the ventral attention network during early adolescence. Negative epicenters were finally situated in the frontal and parietal regions during late adolescence (*p_spin_*< 0.05). Positive disease epicenters in EOS were in the sensorimotor and visual networks during childhood and early adolescence, while spread to widespread regions in late adolescence. Stage-specific structural epicenters were shown in **Figure S8**.

**Tables**

**Table S1.** Demographic and clinical characteristics.

| **Characteristic** | **TD** | **EOS** | **Group comparisons** | |
| --- | --- | --- | --- | --- |
|  |  |  | Statistic values | *p* values |
| ***Overall***  *(7–17 years old)* | *N = 99* | *N = 95* |  |  |
| Sex (male/ female) | 38/ 61 | 36/ 59 | 0.005 ^a^ | 0.94 |
| Age (years) | 14.32 ± 2.08 | 14.61 ± 1.96 | 4381 ^b^ | 0.41 |
| ***Childhood***  *(7–13 years old)* | *N = 39* | *N = 30* |  |  |
| Sex (male/ female) | 16/ 23 | 12/ 18 | 0.007 ^a^ | 0.93 |
| Age (years) | 12.19 ± 1.47 | 12.34 ± 1.59 | 533.5 ^b^ | 0.54 |
| ***Early adolescence***  *(14–15 years old)* | *N = 27* | *N = 39* |  |  |
| Sex (male/ female) | 6/ 21 | 15/ 24 | 1.94 ^a^ | 0.16 |
| Age (years) | 14.89 ± 0.51 | 15.01 ± 0.56 | 453 ^b^ | 0.34 |
| ***Late adolescence***  *(16–17 years old)* | *N = 33* | *N = 26* |  |  |
| Sex (male/ female) | 16/ 17 | 9/ 17 | 1.15 ^a^ | 0.28 |
| Age (years) | 16.38 ± 0.60 | 16.62 ± 0.65 | 307 ^b^ | 0.05 |

*Note: Mean ± SD. TD, typically developing controls; EOS, early-onset schizophrenia patients. ^a^ The χ^2^ value for gender distribution was obtained by chi-square test; ^b^ The U values were obtained by Mann-Whitney test.*

**Table S2. Group differences of cortical thickness in EOS.**

|  | **Brain regions** | **Side** | **Network** | **T values** |
| --- | --- | --- | --- | --- |
| EOS>TD |  |  |  |  |
|  | Visual_15 | L | VIS | 3.32 |
| EOS<TD |  |  |  |  |
|  | Post_2 | L | DAN | −3.88 |
|  | Post_9 | L | DAN | −3.29 |
|  | ParOper_1 | L | VAN | −3.98 |
|  | TempPole_1 | L | LMB | −3.21 |
|  | Temp_1 | L | DMN | −3.61 |
|  | Temp_6 | L | DMN | −2.99 |
|  | PFC_1 | L | DMN | −3.04 |
|  | PFC_10 | L | DMN | −3.42 |
|  | Post_5 | R | DAN | −3.57 |
|  | Post_8 | R | DAN | −3.17 |
|  | Post_9 | R | DAN | −3.63 |
|  | OFC_4 | R | LMB | −3.19 |
|  | pCunPCC_4 | R | DMN | −3.16 |

*Note: VIS, visual network; DAN, dorsal attention network; VAN, ventral attention network; LMB, limbic network; DMN, default mode network; PostC, post central; ParOper, parietal operculum; TempPole, temporal pole; Temp, temporal; PFC, prefrontal cortex; OFC, orbital frontal cortex; pCunPCC, precuneus posterior cingulate cortex.*

**Table S3.** Functional and structural epicenters of EOS.

| **Networks** | **Functional epicenters** | **r values** | **Structural epicenters** | **r values** |
| --- | --- | --- | --- | --- |
| **VIS** |  |  |  |  |
|  | L_Vis_5 | 0.21 | L_Vis_5 | 0.27 |
|  | L_Vis_6 | 0.19 | L_Vis_6 | 0.20 |
|  | L_Vis_9 | 0.22 | L_Vis_9 | 0.26 |
|  | L_Vis_11 | 0.26 | L_Vis_11 | 0.23 |
|  | L_Vis_12 | 0.24 | L_Vis_12 | 0.22 |
|  | L_Vis_14 | 0.21 | L_Vis_14 | 0.18 |
|  | L_Vis_15 | 0.33 | L_Vis_15 | 0.21 |
|  | L_Vis_18 | 0.32 | L_Vis_18 | 0.23 |
|  | L_Vis_19 | 0.24 | L_Vis_19 | 0.21 |
|  | L_Vis_21 | 0.20 | L_Vis_21 | 0.19 |
|  | L_Vis_24 | 0.22 | L_Vis_24 | 0.21 |
|  | R_Vis_4 | 0.16 | R_Vis_4 | 0.14 |
|  | R_Vis_6 | 0.19 | R_Vis_6 | 0.20 |
|  | R_Vis_8 | 0.19 | R_Vis_8 | 0.21 |
|  | R_Vis_9 | 0.17 | — | — |
|  | R_Vis_10 | 0.20 | R_Vis_10 | 0.17 |
|  | R_Vis_11 | 0.23 | R_Vis_11 | 0.14 |
|  | R_Vis_13 | 0.22 | R_Vis_13 | 0.21 |
|  | R_Vis_15 | 0.32 | R_Vis_15 | 0.16 |
|  | R_Vis_18 | 0.23 | R_Vis_18 | 0.15 |
|  | R_Vis_19 | 0.20 | — | — |
|  | R_Vis_21 | 0.27 | R_Vis_21 | 0.16 |
|  | R_Vis_23 | 0.20 | — | — |
|  | — | — | L_Vis_4 | 0.14 |
|  | — | — | L_Vis_10 | 0.20 |
|  | — | — | L_Vis_13 | 0.17 |
|  | — | — | L_Vis_16 | 0.22 |
|  | — | — | L_Vis_22 | 0.18 |
|  | — | — | L_Vis_23 | 0.12 |
|  | — | — | L_Vis_25 | 0.13 |
|  | — | — | L_Vis_26 | 0.16 |
|  | — | — | L_Vis_27 | 0.19 |
|  | — | — | R_Vis_12 | 0.13 |
|  | — | — | R_Vis_20 | 0.16 |
| **SMN** |  |  |  |  |
|  | L_SomMot_19 | 0.15 | — | — |
|  | L_SomMot_20 | 0.16 | — | — |
|  | L_SomMot_22 | 0.17 | — | — |
|  | L_SomMot_28 | 0.16 | — | — |
|  | L_SomMot_35 | 0.16 | — | — |
|  | L_SomMot_36 | 0.17 | — | — |
|  | R_SomMot_25 | 0.15 | — | — |
|  | R_SomMot_26 | 0.15 | — | — |
|  | R_SomMot_29 | 0.16 | — | — |
|  | R_SomMot_34 | 0.18 | — | — |
|  | R_SomMot_35 | 0.15 | — | — |
|  | R_SomMot_40 | 0.15 | — | — |
| **DAN** |  |  |  |  |
|  | L_Post_9 | −0.17 | — | — |
|  | R_Post_8 | −0.18 | — | — |
|  | R_Post_9 | −0.15 | — | — |
|  | — | — | L_Post_3 | −0.11 |
|  | — | — | R_Post_6 | −0.12 |
| **VAN** |  |  |  |  |
|  | L_ParOper_2 | −0.15 | — | — |
|  | L_ParOper_4 | −0.15 | — | — |
|  | L_FrOperIns_3 | −0.14 | — | — |
|  | R_TempOccPar_7 | −0.17 | R_TempOccPar_7 | −0.16 |
|  | — | — | L_FrOperIns_7 | −0.12 |
|  | — | — | R_TempOccPar_3 | −0.12 |
|  | — | — | R_TempOccPar_6 | −0.14 |
| **LMB** |  |  |  |  |
|  | R_OFC_4 | −0.12 | — | — |
| **FPN** |  |  |  |  |
|  | L_Par_2 | −0.21 | — | — |
|  | L_Par_3 | −0.16 | — | — |
|  | L_Par_5 | −0.13 | — | — |
|  | L_Par_6 | −0.19 | — | — |
|  | L_PFCv_1 | −0.15 | — | — |
|  | R_Par_2 | −0.14 | R_Par_2 | −0.13 |
|  | R_Par_4 | −0.19 | R_Par_4 | −0.14 |
|  | R_Par_5 | −0.13 | R_Par_5 | −0.11 |
|  | R_PFCv_1 | −0.16 | — | — |
|  | R_PFCl_2 | −0.12 | — | — |
|  | R_PFCl_4 | −0.13 | — | — |
|  | — | — | L_Temp_1 | −0.12 |
|  | — | — | R_Par_1 | −0.14 |
|  | — | — | R_Par_3 | −0.13 |
| **DMN** |  |  |  |  |
|  | L_Temp_1 | −0.14 |  |  |
|  | L_Temp_4 | −0.13 |  |  |
|  | L_Temp_7 | −0.15 |  |  |
|  | L_Temp_10 | −0.11 |  |  |
|  | L_PFC_1 | −0.15 | — | — |
|  | L_PFC_2 | −0.12 | — | — |
|  | L_PFC_5 | −0.17 | — | — |
|  | L_PFC_7 | −0.15 | — | — |
|  | L_PFC_10 | −0.15 | — | — |
|  | R_Par_4 | −0.13 | — | — |
|  | R_Temp_5 | −0.12 | — | — |
|  | R_Temp_6 | −0.12 | — | — |
|  | R_PFCv_1 | −0.14 | — | — |
|  | R_PFCv_2 | −0.15 | — | — |
|  | R_PFCv_3 | −0.16 | — | — |
|  | — | — | L_Temp_3 | −0.11 |
|  | — | — | L_PFC_23 | −0.11 |
|  | — | — | L_PFC_24 | −0.12 |
|  | — | — | R_Par_5 | −0.13 |

***Note:*** *SMN, sensorimotor network; FPN, frontoparietal control network; FrOper, frontal operculum; TempOcc, temporal occipital; Par, parietal; PFCv, ventral prefrontal cortex; PFCl, lateral prefrontal cortex.*

**Figures**

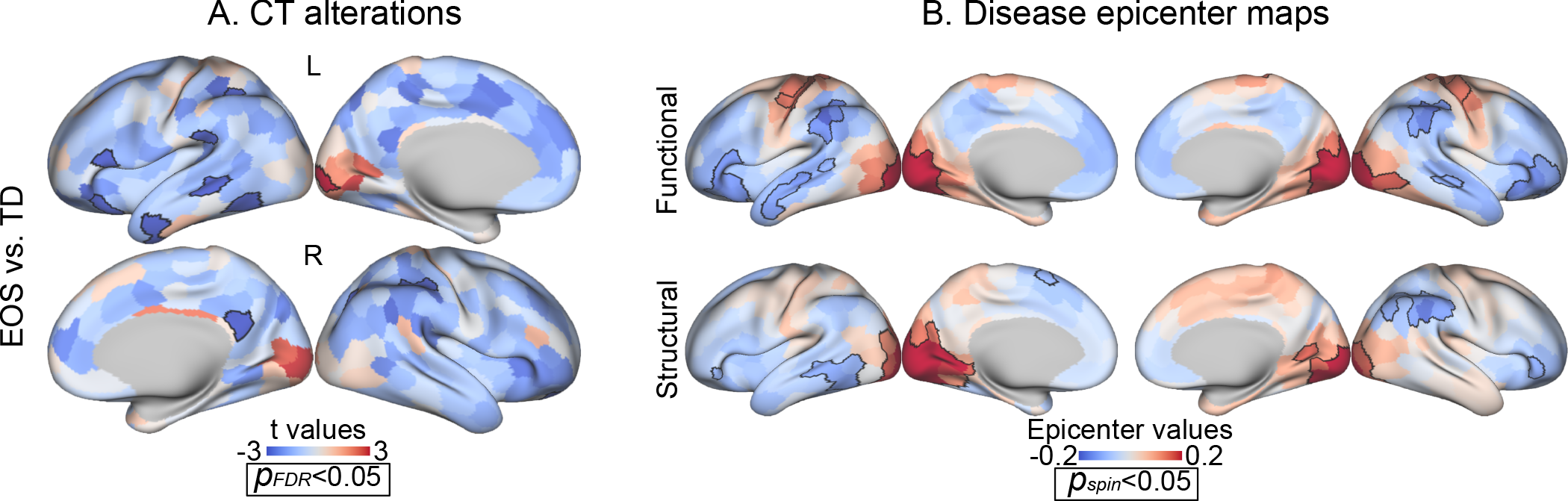

**Figure S1.** Replicated disease epicenters with squared age as one of covariates.

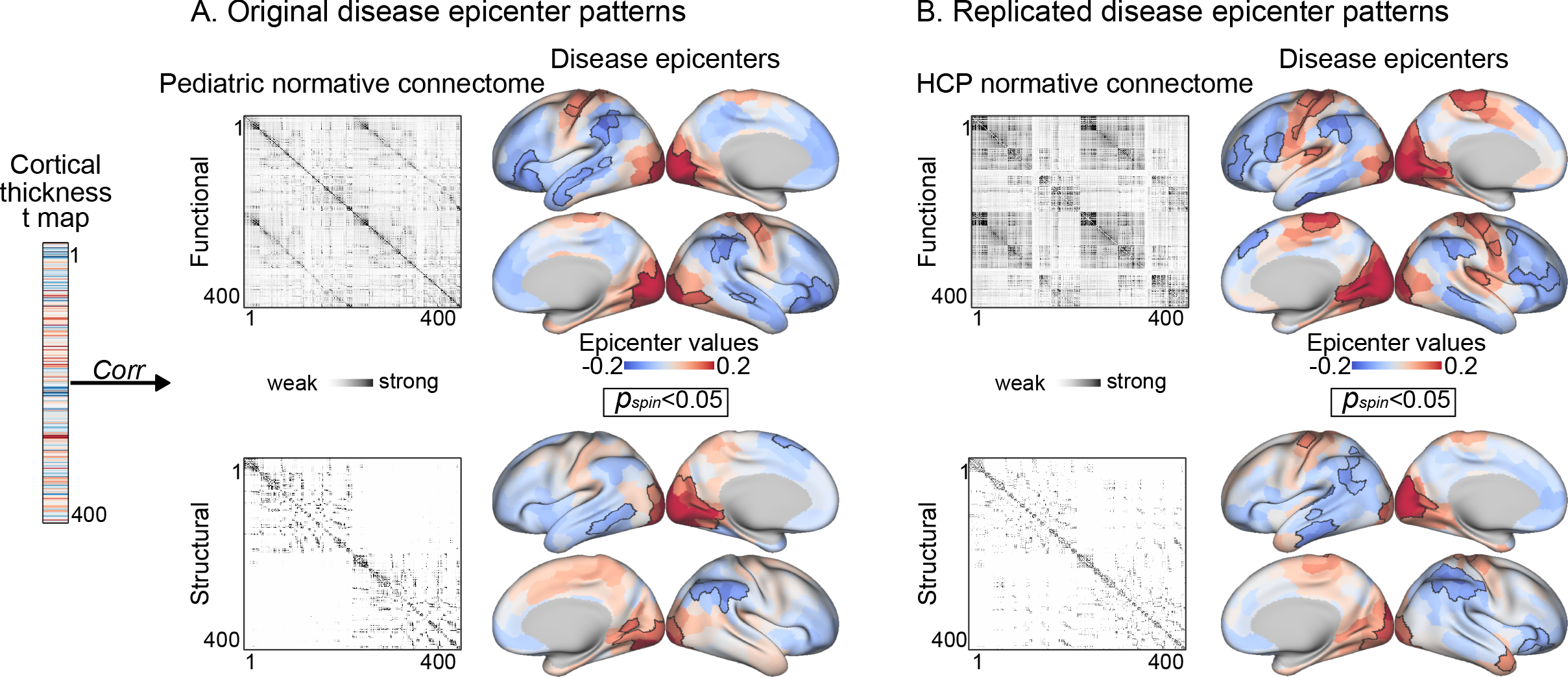

**Figure S2.** Replicated disease epicenters by using normative adult connectome derived from HCP data.

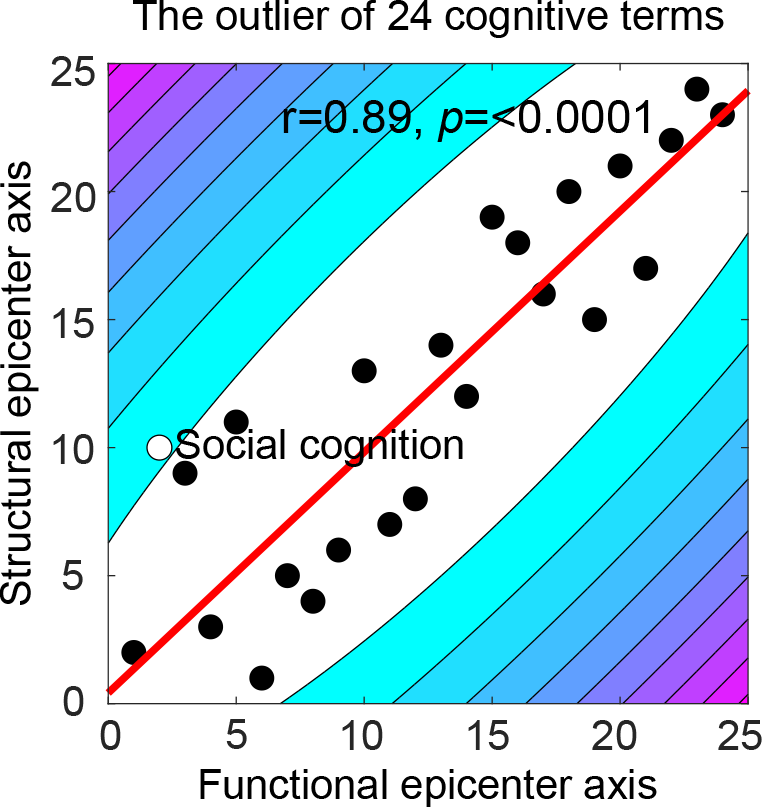

**Figure S3.** The outlier of 24 cognitive terms. The outlier was identified by means of bootstrapped Mahalanobis distance . After removing the outlier, Shepherd’s pi correlation was then computed between functional and structural ranked cognitive terms (*r* = 0.89, *p* < 0.0001).

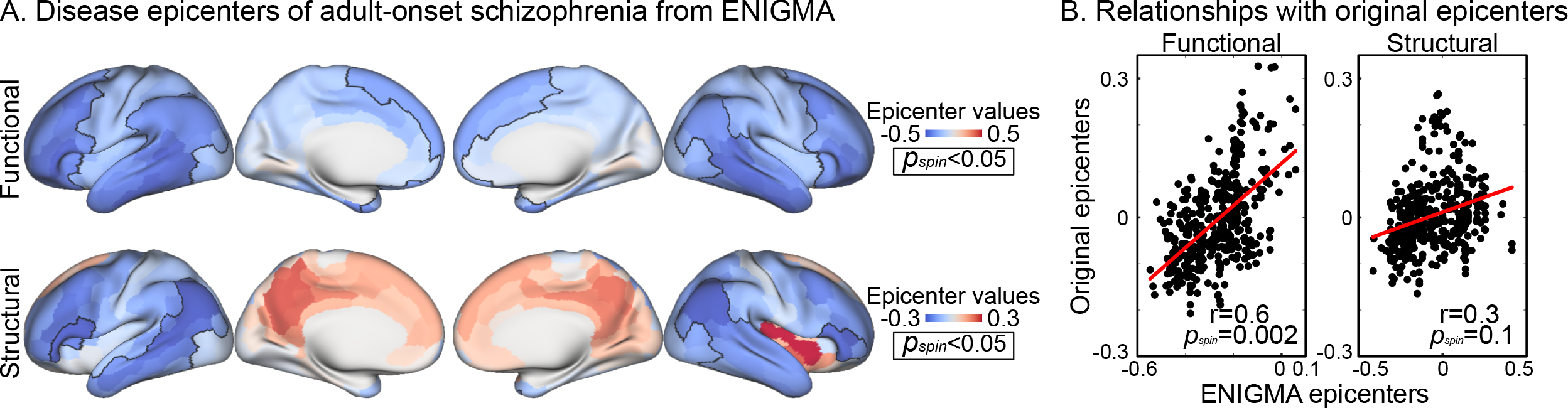

**Figure S4.** Disease epicenters of adult-onset schizophrenia from the ENIGMA Toolbox.

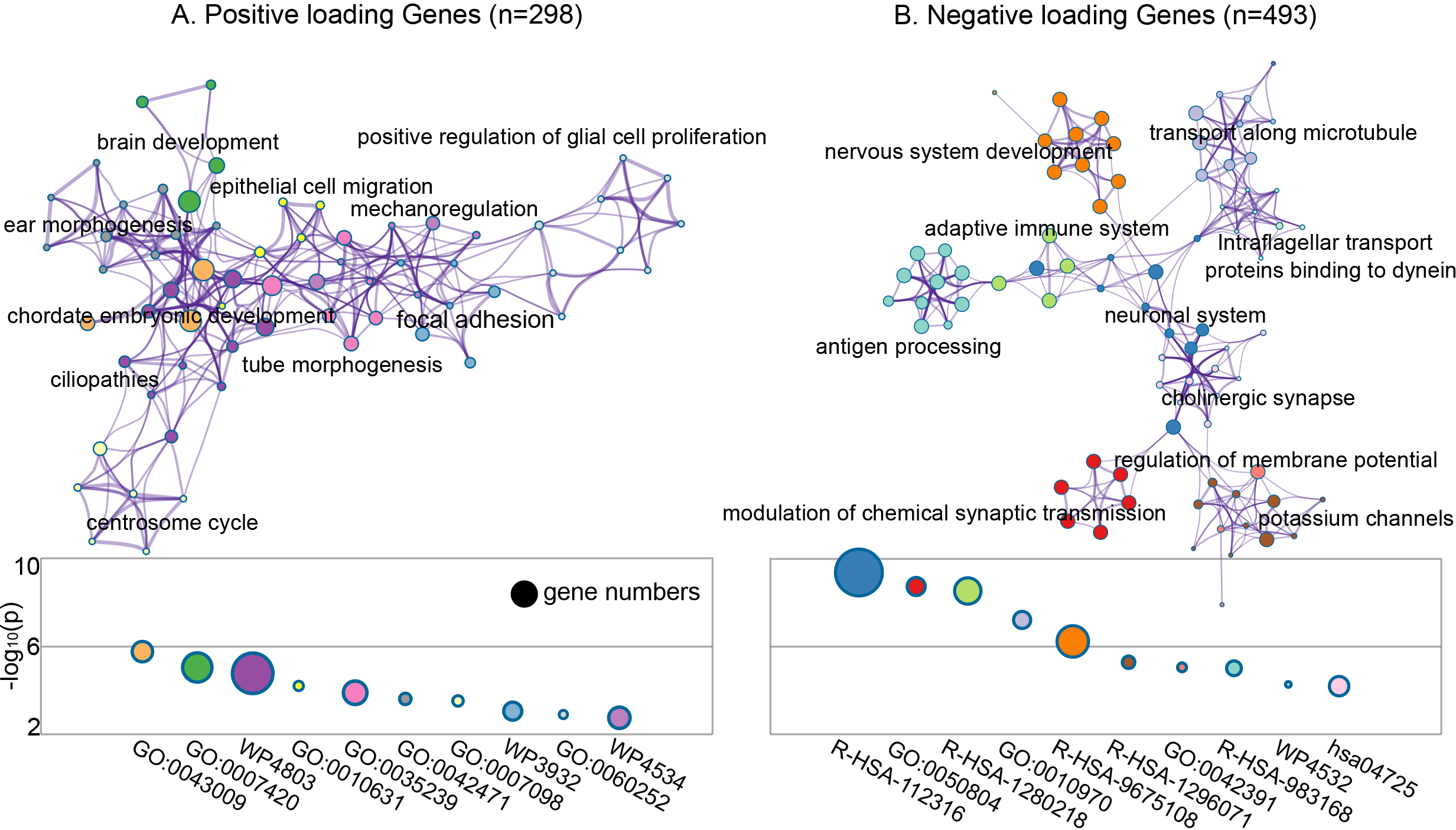

**Figure S5.** Underlying biological processes. Metascape enrichment network visualization was performed for PLS+ and PLS− gene set. A circle node refers to an enriched term (the same color reflects the same cluster) and size of the node is proportional to the number of included genes.

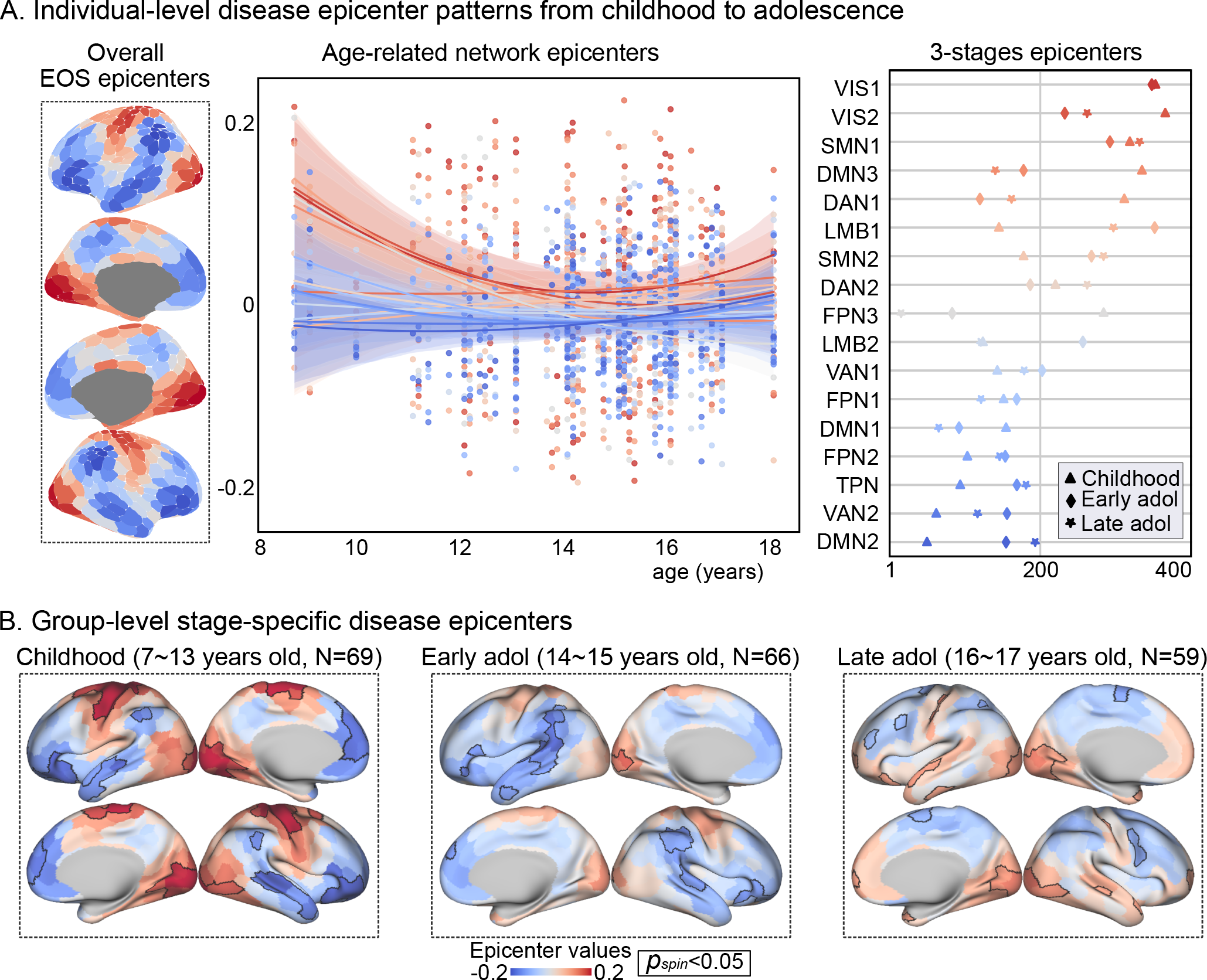

**Figure S6.** Functional disease epicenter dynamics as a function of age of onset.

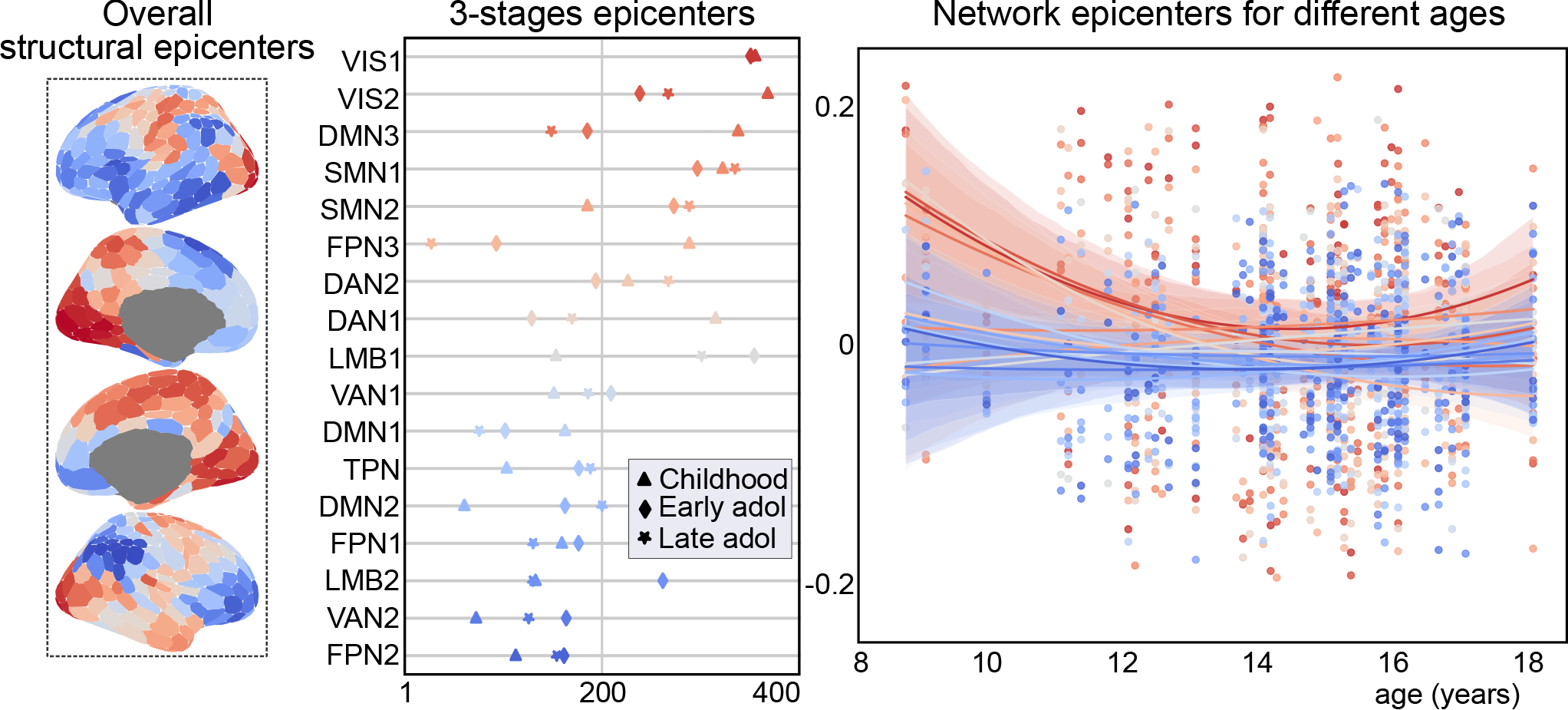

**Figure S7.** Structural disease epicenter dynamics generated by individual-level epicenters.

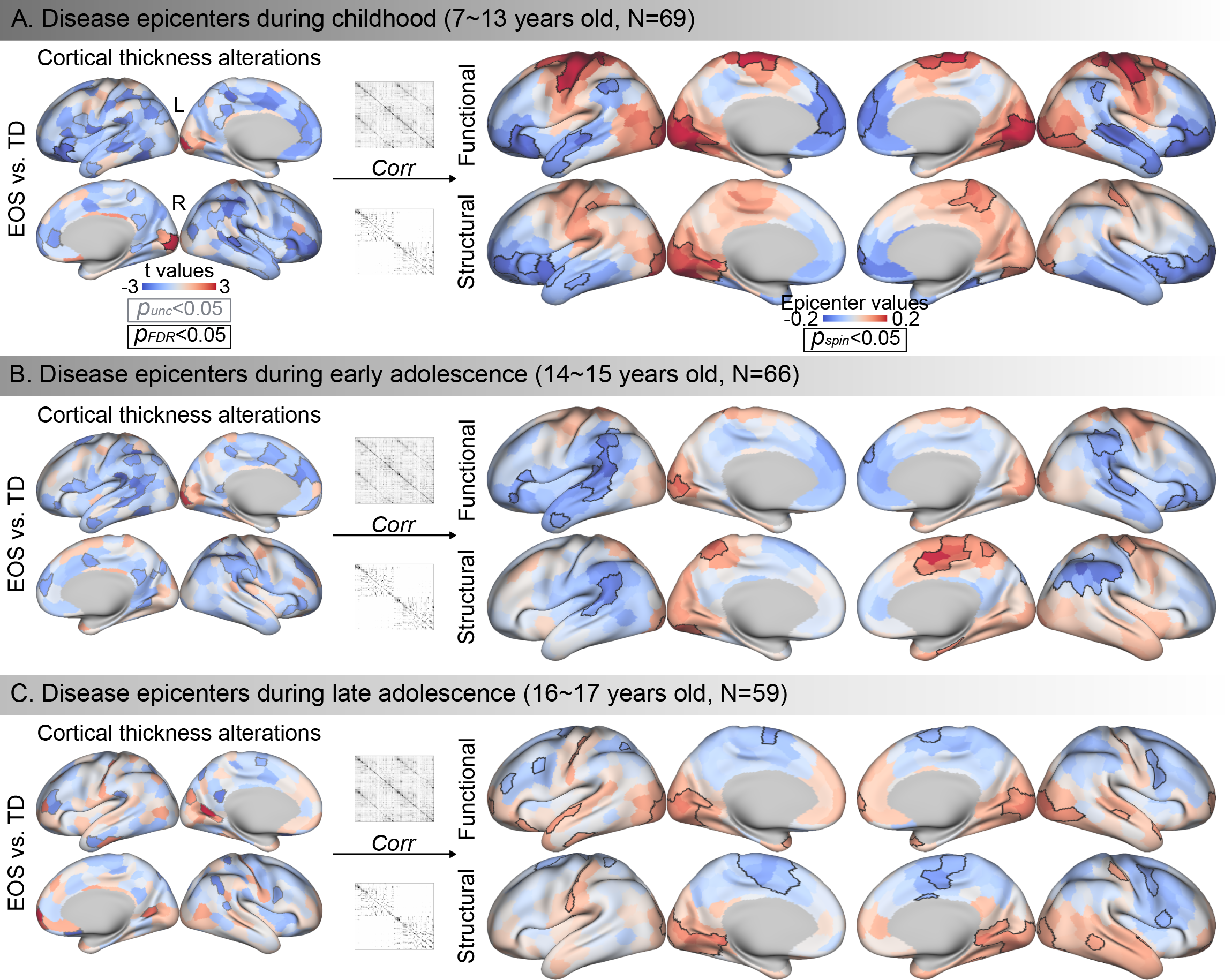

**Figure S8.** Functional and structural epicenter dynamics generated by group-level epicenters.
